# Biosys-LiDeOGraM: A visual analytics framework for interactive modelling of multiscale biosystems

**DOI:** 10.1101/2023.06.23.546209

**Authors:** Nathalie Mejean Perrot, Severine Layec, Alberto Tonda, Nadia Boukhelifa, Fernanda Fonseca, Evelyne Lutton

**Affiliations:** Université Paris-Saclay, INRAE, AgroParisTech, UMR 518 MIA Paris-Saclay, F-91120 Palaiseau, France; Université Paris-Saclay, INRAE, AgroParistech, Micalis Institute, UMR1319, F-78350 Jouy-en-Josas, France; ISCPIF, F-75013 Paris, France; Université Paris-Saclay, INRAE, AgroParisTech, UMR SayFood, F-91120 Palaiseau, France

**Keywords:** Multi-scale modelling, interactive design, multi-data sets, integrative biology, computational biology, visualization, machine learning, visual analytics, Lactic Acid Bacteria

## Abstract

In this paper, we present a test of an interactive modelling scheme in real conditions. The aim is to use this scheme to identify the physiological responses of microorganisms at different scales in a real industrial application context. The originality of the proposed tool, Biosys-LiDeOGraM, is to generate through a human–machine cooperation a consistent and concise model from molecules to microbial population scales: If multi-omics measurements can be connected relatively easily to the response of the biological system at the molecular scale, connecting them to the macroscopic level of the biosystem remains a difficult task, where human knowledge plays a crucial role. The use-case considered here pertains to an engineering process of freeze-drying and storage of Lactic Acid Bacteria. Producing a satisfying model of this process is a challenge due to (i) the scarcity and variability of the experimental dataset, (ii) the complexity and multi-scale nature of biological phenomena, and (iii) the wide knowledge about the biological mechanisms involved in this process. The Biosys-LiDeOGraM tool has two main components that can have to be utilized in an iterative manner: the Genomic Interactive Clustering (GIC) module and the Interactive Multi-Scale modellIng Exploration (IMSIE) module, both involve users in their learning loops. Applying our approach to a dataset of 2,741 genes, an initial model, as a graph involving 33 variables and 165 equations, was first built. Then the system was able to interactively improve a synthetic version of this model using only 27 variables and 16 equations. The final graph providing a consistent and explainable biological model. This graphical representation allows various user interpretations at local and global scales, an easy confrontation with data, and an exploration of various assumptions. Finally Biosys-LiDeOGraM is easily transferable to other use-cases of multi-scale modelling using ‘functional’ graphs.

**Author summary:** The use of “omics” data for understanding biological systems has become prevalent in several research domains. However, the data generated from diverse macroscopic scales used for this purpose is highly heterogeneous and challenging to integrate. Yet, it is crucial to incorporate this information to gain a comprehensive understanding of the underlying biological system. Although various integrative analysis methods that have been developed provide predictive molecular-scale models, they only offer a mechanistic view of the biological system at the cellular level. In addition, they often focus on specific biological hypotheses through dedicated case studies, making it difficult to apply their results to other scientific problems. To address these issues, we propose an interactive multi-scale modelling approach to integrate cross-scale relationships providing predictive and potentially explanatory models. A proof-of-concept tool has been developed and was validated in the context of the bioproduction of *Lactococcus lactis*, a bacterial species of high economic interest in the food industry and for which the control of the bioprocess is essential to guarantee its viability and functionality. Our approach can be applied to any biological system that can be defined through a set of variables, constraints and scales.

## Introduction

Over the past two decades, knowledge management in biological research has experienced a profound transformation as a result of the availability of high-throughput experimental technologies and advances in mathematical and computer modelling. Consequently, large omics datasets are now commonly found in studies focused on integrated, system-level approaches in biology [1, 2]. The interpretation of omics data is also largely enabled by the availability of multiple reference databases and several open-source bioinformatics software packages. These packages allow the combination of different types of high-dimensional data at the genome level, and thus help experts reach new insights on physiological responses, from genotype to phenotype [3, 4].

Computer modelling and machine learning have also become increasingly important for the integration of multi-omics datasets to address fundamental biological questions [5]. These approaches have enabled the establishment of biological networks and have facilitated the extension of analyses to a more complete, multi-scale reconstruction of biological processes, thereby providing comprehensive overviews of entire biological systems [1, 6]. The integration of these multi-omics datasets has thus led to major advances in the holistic understanding of the molecular mechanisms that govern the functional dynamics of living individual cells and their communities in response to environmental change [2, 7, 8].

Typically, sophisticated computational algorithms are required to integrate large omics datasets in order to facilitate their reuse and connection with other data resources. Some modelling approaches are widely used for the analysis of biological systems [1, 2, 7, 8]. These approaches are mainly deterministic and based on differential equations, established through known biological events and relationships. They have been applied, for instance, to describe dynamic molecular events at a cellular scale in the case of a homogeneous biological system [9]. However, the paucity of informative scientific data and related measurements, as well as the high levels of noise and uncertainty in regulatory networks, are limiting factors which often lead to aberrant results [3]. Deterministic systems are therefore considered unsuitable for the analysis of population dynamics and for the investigation of complex multi-scale biological systems [10].

Since biological systems are always subject to unknown effects that can occur at any scale, from molecular to macroscopic, stochastic models have been applied. They are now widely used to provide a clear and accurate overview of molecular mechanisms in spatially heterogeneous dynamical systems [11, 12]. Although more powerful and accurate, these methods have limitations. Statistical approaches generally need large sets of physiological parameters that are often not easily accessible through experiments. Moreover, these statistical approaches rely primarily on *a priori* knowledge to reduce the scope of the investigation, thus limiting the ability to identify unknown behavior [13]. In addition, these methods frequently struggle to incorporate high dimensional datasets that are both large and complex in terms of size, heterogeneity, and content [14].

Usually, Bayesian statistical or frequentist inference methods are the most widely used approaches for modelling complex multiscale, multi-level and multi-dimensional aspects of biological systems [15] These mathematical approaches have the advantage of offering a better prediction of the behavior of individuals facing environmental change, based on various characteristics that are known or assumed to be associated with some scientific evidence [1]. They are mainly used to identify all interactions between molecular components in order to build a cell network model that facilitates simulation and analysis [1]. Nowadays, Bayesian inference methods are widely used in systems biology and are essentially applied in the field of clinical diagnosis in human health and medicine. The goal is to provide a better understanding of individual cells or cell population physiological response to disease through a model of spatio-temporal dynamics and structure at the molecular level [3, 15]. However, these multi-scale mathematical models are usually tailored to a particular multicellular system, and created to address a specific biological issue, under specific time kinetics and controlled experimental conditions. On the one hand, this leads to a limitation in their scope of application since these models cannot be easily extrapolated to other biological systems due to their high complexity, their purely data-driven approach and in some cases, the lack of reference data [15]. On the other hand, it remains difficult to achieve a realistic integration of large-scale metadata sets resulting from increasingly sophisticated questions and experiments, while taking into account all steps of the bioengineering process. This is a real challenge for computational multi-scale modelling [14, 16, 17].

Nowadays, the knowledge in integrative biology is still poorly characterized. Some authors propose some guidelines for computing approaches [3, 18], and promote graph theory as a good candidate for multi-scale reconstruction, though it has the potential for error propagation if there are errors in the graph structure. Indeed, the emphasis is on the initialization of links between variables, with a priori knowledge provided by experts in biology [3, 19]. [20] shows that the most robust predictive models are created by coupling machine learning and multi-scale modelling.

In current multi-scale modelling methods, visualization is often used to represent biological knowledge as well as statistical findings from machine learning methods. However, it typically provides limited opportunities for biological experts to participate in the learning process, which hinders integrative biological research [3, 19]. Indeed, the integration of human knowledge and expertise into machine learning is becoming increasingly popular across various domains and has proven to be effective [21]. Thus, another challenge for integrative biology is to engage domain experts in the different steps of the modelling process, for example, by implementing semi-automatic iterative learning loops.

In this paper, we propose a Visual Analytics (VA) framework (Fig. 1), which allows biological experts to construct and evaluate various computational models, based on a combination of their own knowledge with available multi-scale data. VA is defined as the science of analytical reasoning facilitated by interactive visual interfaces [22]. More generally, the goal of VA is to support semi-automatic analytical processes, by combining human and computational capabilities to achieve the most effective results [23].

**Fig 1.**
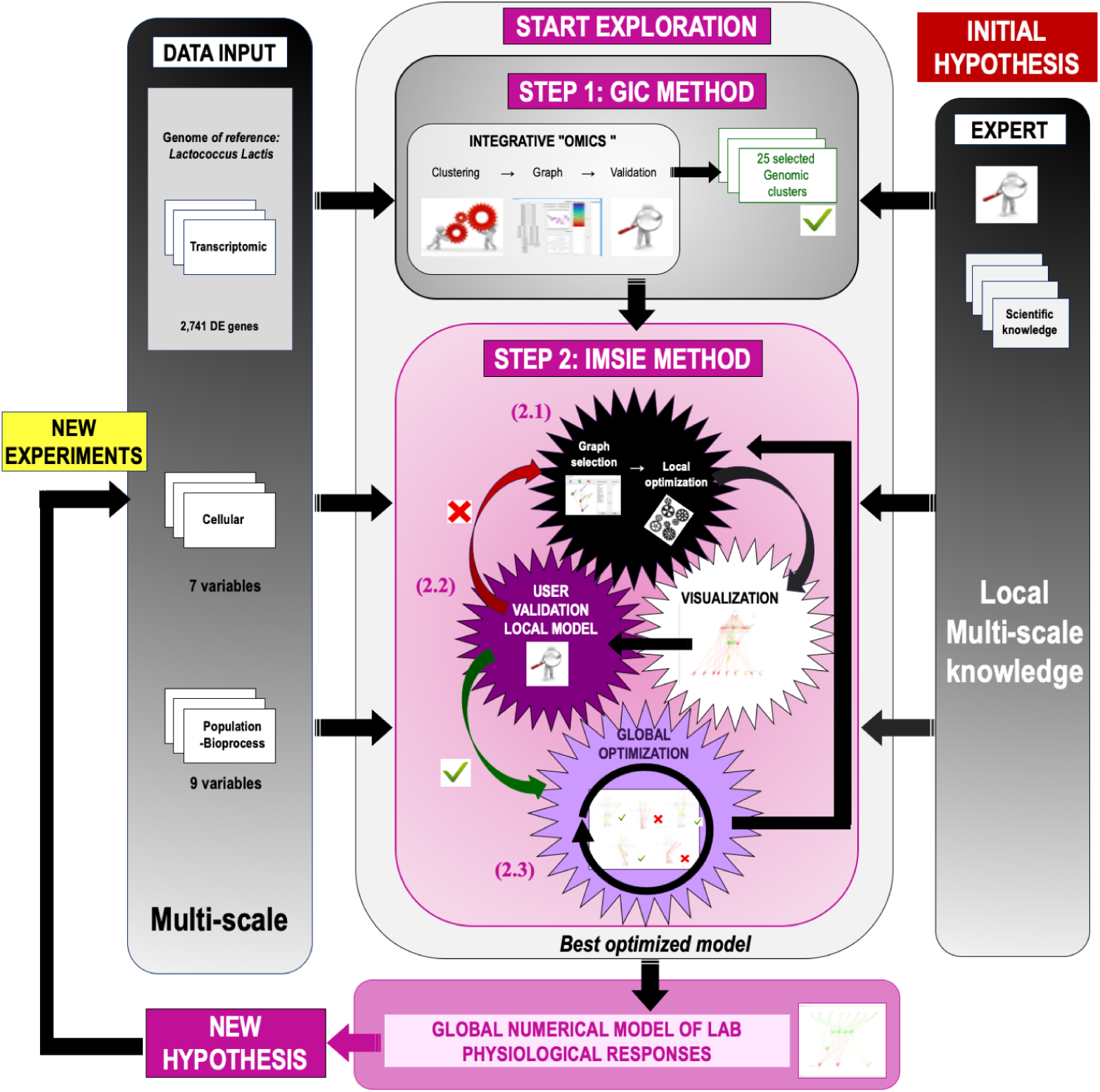
Overview of the visual analytics workflow. *Right:* Domain experts come to the experiment with initial hypotheses and local knowledge. *Left:* Transcriptomic and genomic data from a biological experiment consisting of variables at four distinct scales: genome, cell, population and bioprocess. *Middle:* The proposed framework composed of two steps. In STEP 1 (GIC method), domain experts visualize the various clusters of variables extracted by machine learning (hierarchical clustering) and select those that make sense to them. In our use-case, a total of 25 gene clusters were chosen. In STEP 2 (IMSIE method), experts visually examine a set of local models (i.e. a set of possible mathematical equations linking variables at different scales). In this framework, explicit knowledge is manifested as constraints on the graph (step 2.1), while implicit expert knowledge is leveraged through the evaluation and selection of local models (step 2.2). If this selection provides good predictions, domain experts can trigger step 2.3. If the prediction is judged poor, then steps 2.1 and 2.2 can be revisited resulting in new sets of constraints and equations. Step 2.3 optimizes a global model as a combination of the local models selected by experts. All steps are iterated until a satisfactory global model is found.

As a proof of concept, this framework was applied to a multi-scale engineering process involving *Lactococcus Lactis* Lactic Acid Bacteria (LAB), commonly used in food industrial bio-transformation and recognized for their health benefits as probiotics [24]. Regardless of their industrial applications, these bacteria must be able to withstand sometimes drastic stress conditions during their production processes (fermentation and stabilization via freezing or freeze-drying) that can affect the cellular physiology [25, 26], influence their enzymatic activities, decrease their stability and lead to loss of viability [27, 28]. Although nowadays, several studies have determined various ways to improve their resistance [25, 29–31], bacterial survival is highly dependent on the cell physiology of LAB confronted with stresses at each step of the bioprocess. In order to validate this framework, we rely on a combination of multi-scale datasets including the transcriptome, cell membrane fatty acid composition, biological activity (acidifying activity and viability) following different fermentation conditions (culture temperature and cell harvesting time) that have been generated in a spatio-temporal biological context along a bioprocess chain including the fermentation, stabilisation and storage steps. The objective is to be able to provide the biological expert with new perspectives for the optimization of LAB bioprocess engineering.

## Materials and methods

### Use-case: viability of LAB in a freeze-drying industrial process

Many studies have shown that the robustness of LAB cells in industrial processes is dependent on

1. the cell growth conditions, medium composition [32, 33], fermentation temperature [25, 34–36], fermentation pH [33–37], and cell harvesting time [25, 33, 38].
2. the physico-chemical factors of stabilization, such as freezing and freeze-drying parameters [31, 39, 40], use of lyoprotectants [26, 41–44], storage [31, 32, 45, 46] and rehydration conditions [47–49].

In addition, transcriptome studies provide insight into a microorganism’s overall response to stress, often leading to survival under normally lethal conditions [24, 50].

Therefore, the biological data used in this project were derived from several experimental approaches on *Lactoccocus lactis* subsp. *lactis* TOMSC161, hereafter called *L. lactis* TOMSC161, a strain of industrial interest that is robust to the stressful conditions that occur during fermentation and stabilization processes, the results of which were previously published by [25, 51]. In summary, the described datasets consist of: (i) the colony-forming units (UFC) counts;(ii) acidifiying activity measurements; (iii) fatty acid data obtained in GC-HPLC; (iv) anisotropies, and (v) transcriptomic data from the study of cells harvested at an early stationary growth-phase (ES) and a late stationary growth-phase (LS) (6h after ES) during two independent fermentations leading to 22 °C and 30 °C, and repeated in triplicate [25]. The objective of these works was to get a deep understanding of the effect of each step of the fermentation, stabilization and storage processes that could influence the biological system, affect their viability and influence the behavior of LAB. This would enable the preservation of their functional properties by choosing the best control parameters that could be used throughout the production chain.

### Data inputs of Biosys-LiDeOGraM: a multi-scale description of the biosystem

The cell concentrates were characterized at each step of the stabilization process: concentration, lyoprotectants addition, freezing, freeze-drying, and after three months of storage. In addition, transcriptomic data in early and late stationnary growth phases of fermentation at 22 and 30 °C have been included in the dataset. In total, 16 variables at different mesoscopic scales and 2,741 differentially expressed (DE) genes were considered. The various measured variables are related to four different scales as follows (Table 1).

**Table 1.**
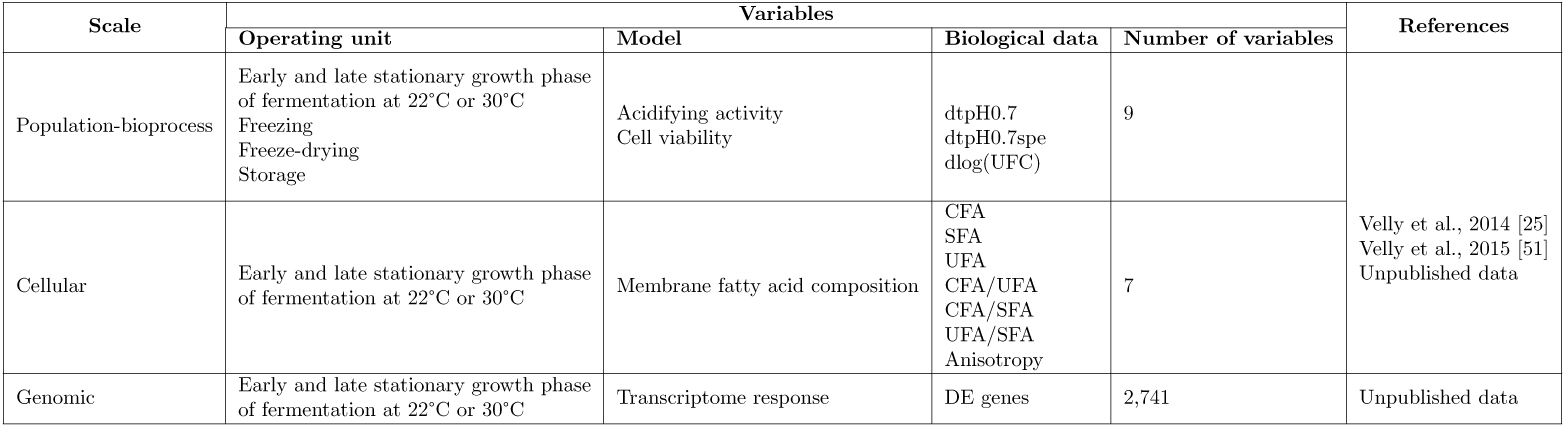
Summary of variables considered in the use-case.

### Acidifying activities at the bioprocess scale

Starters LAB strains are usually marketed on the basis of their acidifying efficiency, which is essential in the industrial manufacturing processes of fermented products [52]. Therefore, the acidifying activity dataset was also included in our multi-scale application. Measurements of the acidifying activity of this strain were carried out on cells harvested at 3 h from the stationary growth phase of fermentation, and were obtained continuously throughout the bacteria growth by the Cinac system, as described in the previous work [25]. The indicator *tpH*07 (corresponding to the time required to obtain a decrease of 0.7 pH units, in min) was used to characterize the acidifying activity of cells, in order to determine the effect of each step of the stabilization process (concentration, lyoprotectants addition, freezing, freeze-drying, and after one or three months of storage). In addition, the specific acidifying activity (named *tpH0.7spe*), in between the bioprocess and population scales, taking into account both the cell viability and the acidifying activity [51] was also considered. The indicator *tpH0.7spe* (corresponding to the ratio of *tpH*0.7*/log*(*UFC*), with UFC corresponding to colony-forming units (see below), and expressed in min (log (UFC/mL))*^−^*^1^ ensures a global biological activity of starters LAB. Loss of acidification activity was determined at each step of the process as follows [25, 51]:

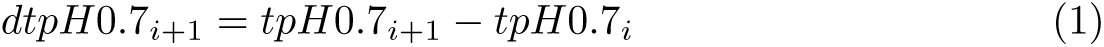

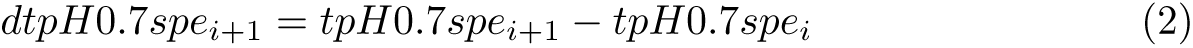

where *i* and *i* + 1 are two subsequent steps of the stabilization process.

### Cell viability at the population scale

The effect of freeze-drying and storage processes on cell viability was identified in a previous study [25] and considered in multi-scale integration. The cell viability was measured at each step of the stabilization process, by counting UFC per mL after 48 h incubation at 30°C and anaerobiosis. Loss of cell viability was evaluated at each step of the process according to the following equation as mentioned in [25]:

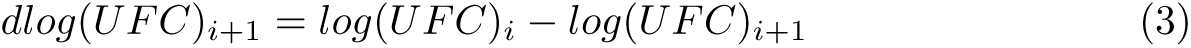

where *i* and *i* + 1 are again two subsequent steps of the process.

### Cell membrane at the cellular scale

The cell envelope of microorganisms plays a crucial role in cellular integrity. Specifically, the survival of bacteria depends on their ability to maintain phospholipid membrane properties to meet environmental requirements. However, the cell membrane has been reported as a major site of damage during freezing, freeze-drying or rehydration where transitions of membrane phases occur [30]. This results in the modification of membrane structure, viscosity and permeability, leading to viability and acidifying activity looses [53]. Therefore, the cell membrane properties of *L. lactis* TOMSC161 for each tested condition were integrated in this analysis. The biological datasets used to characterize cell membrane modifications have already been published in [51]. These include measurements of: (i) fatty acid composition (i.e. saturated fatty acids (SFA), unsaturated fatty acids (UFA) and cyclic fatty acids (CFA)); and (ii) fluorescence anisotropy, which provides information on the level of membrane fluidity. In addition, the following fatty acid ratios UFA/SFA, CFA/UFA and CFA/SFA are generally calculated to interpret the impact of freeze-drying and storage on cell membrane composition, and therefore taken into account in the equations.

### Transcriptome responses at the genomic scale

Many studies on LAB were also referenced to understand the molecular mechanisms that govern an adaptive response in these microorganisms, leading to a physiological state of tolerance, and thus survival under normally lethal stress conditions [24, 50]. In previous work, the physiological responses of *L. lactis* TOMSC161 have been investigated by RNA-Seq transcriptome analysis to understand its tolerance to freeze-drying and storage (data not published). RNA-Seq data used in this study were deposited in the Gene expression Omnibus of the NCBI under the accession number GSE74554. The detailed experimental approaches for RNA extraction, RNA-Seq librairies preparation, RNA-seq sequencing and bioinformatics data analysis were performed as described in [54, 55]. Briefly, only uniquely mapped reads of *L. lactis* TOMSC161 (NCBI Bioproject accession number: PRJEB4823, [56]) were used for data analysis. Differential gene expression was determined by comparing the number of uniquely mapped reads for each gene in the three replicates, from the early and the late stationary growth phases. A count table was generated to select only data corresponding to CDS (CoDing Sequence) features that exhibited an average of more than 50 mapped reads. A differential gene expression analysis between the two conditions was performed using DESeq2 (version 1.8.1) package in the statistical environment R [57]. The *p-values* were adjusted using the Benjamini and Hochberg method, which assesses the false discovery rate [58]. Therefore, overall responses defined by DE genes were also included. Moreover, from transcriptomic data, DE genes were grouped according to their expression profiles and identified as either belonging to the same metabolic pathway gene clusters, or the same operon structure encoding a biological function in the *L. lactis* TOMSC161 genome.

## Methods

### Step 1: Genomic Interactive Clustering (GIC)

One objective of the **Biosys-LiDeOGraM** approach is to reduce the number of variables at the genomic scale while improving prediction performance. Additionally, selecting the most relevant variables [59], i.e. performing a dimensionality reduction, facilitates the modelling process and possibly reduces the associated computational time.

The GIC step is based on a user-guided hierarchical clustering. Among the variety of algorithms dedicated to the analysis of gene expression in the field of bioinformatics [60], hierarchical clustering provides clusters organized in hierarchies, where each cluster is a subset of the next higher cluster in the hierarchy. It is often convenient for visualizing the structure of a dataset and identifying relationships or outliers in the data.

However, a challenge lies in fine-tuning these algorithms. For instance, the ideal number of levels and clusters is rather difficult to estimate, and even when routines to automatically estimate it are available, manual settings are often preferred. Various metrics can also be used to evaluate the distances between gene expression profiles. They need to be appropriately chosen, but efficient parameter tuning is strongly dependant on the application and data [61]. A way to overcome this difficulty is to use a semi-automatic approach, where domain experts can review gene clusters and interactively adjust the parameters.

Another difficulty is due to the small size of the datasets, and the presence of error and noise: proposed clusters could be due to random correlations. To deal with such aberrant clusters, we use a hierarchical density-based spatial clustering of applications with noise (DB-SCAN) method [62–65] (with an *α* parameter set to 0.5): clusters are displayed as a dendrogram, where parent clusters include all elements of child clusters, including elements considered as outliers in child clusters. This decomposition is constrained by a minimum number of elements set by the user.

We choose to group DE genes that have similar expression levels according to the experimental conditions, as similar expression levels can often be associated with the same biological function and/or metabolic pathway. A prior normalization of the gene expression data is used (equation 4),

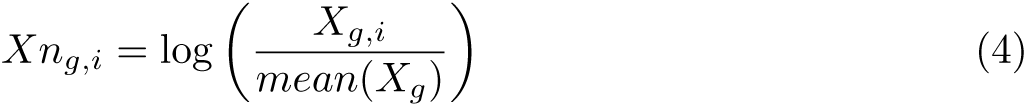

where *X_g_*is a vector made of the expression levels of gene *g* over all experiments, *X_g,i_* is the expression level for gene *g* and experiment *i* and *Xn_g,i_*is the normalized expression level for gene *g* and experiment *i*.

The logarithmic normalization of equation 4 is convenient for fixing a clustering threshold parameter that can distinguish between the average expression value, and the expression levels that are twice as low or twice as high.

We standardize the gene expression by transforming *Xn_g,i_* into a Z-score according to [66]. This involves transforming *Xn_g,i_* so that the average expression of each gene is equal to 0 and the standard deviation is equal to 1. In this way the focus of the clustering is set on the change in expression level between different experiments, regardless of the change in expression level (i.e. fold change) of each gene, which can be different. Finally, it enables us to better distinguish genes whose expression level changes only slightly compared to their average expression level.

### Step 2: Interactive Multi-Scale Modelling Exploration (IMSIE)

Local and global models are built in the IMSIE step via an interactive process. Local models correspond to simple models that represent possible links between a variable and a set of connected variables (parent-variables). Each set of dependencies are considered independently and are represented as equations. A global model is then computed as a coherent chain of local models, according to the architecture and constraints of a graph. This step is based on an approach called LiDeOGraM [67, 68]. Local models are computed using multi-linear regression. Then a global model is optimized with an evolutionary algorithm as follows.

At all steps dependencies are shown to the user as a graph, *G*, where each node corresponds to a variable of the system and each edge is a possible link (for local models) or a proposed link (for global models) between variables. Additionally *G* is displayed with variables organized in rows corresponding to the various scales.

### Least squares multi-linear regression

Orthogonal matching pursuit [69] is a common algorithm for multi-linear regression. In biosystems applications, however, the low amount of available training data can easily lead to overfitting. In order to tackle this issue, we modified the algorithm so that it returns a set of equations of variable complexity, following the general idea that showing different compromises between complexity and fitting to the user might lead to optimal choices.

For each variable *v*, the set of its possible antecedents *A_v_*is extracted from the graph *G*. This set thus includes all variables at lower scales that can explain the value of *v*; its cardinality is noted as *A* = *|A_v_|*.

A sequence of *A* multi-linear regressions (*i* = 1, 2*,…, A*) is then performed, for exploring the set of all possible subsets of variables in *A_v_* ordered according to their size: At iteration *i*, we only consider the subsets of variables *{S_i_*_1_*, S_i_*_2_*,…, S_iK_}* of cardinality *i*. In this way, *i* controls the complexity of the learned equations. Then, a list of *A* equations ordered by complexity is presented to the user.

### Pearson correlation

An equation is a *local model*, it is evaluated according to two metrics, corresponding to: (i) its complexity, defined as the number *i* of terms used in the equation; and (ii) its precision, i.e. a numerical evaluation of how it fits the available data. We use the Pearson correlation coefficient [70] between measured and predicted values.

### Evolutionary optimized global model

A *global model* is then searched as a chain of *local models*, one local model for each variable (selected from the ordered list described above), in a way that is coherent. Finding such a coherent combination is a complex optimization problem.

This problem is combinatorial, it is tractable with a manual or exhaustive search for small instances. However, biosystem datasets typically feature a considerable number of variables, so even if an expert could in theory manually select an equation/local model for each, it is intractable in practice.

IMSIE computes an optimal global model, in the sense that it is able to fit the values of experimental data. For that purpose, we use an evolutionary algorithm [71], a class of stochastic optimization techniques loosely inspired by the neo-Darwinian paradigm of natural selection, able to return solutions of reasonable quality in a limited amount of time.

Once a global model has been computed, the user has access to all equations through an interface and can modify the constraints on the graph, enforce some equations or links, and prohibit some others. All of the above steps can be iterated as much as needed, and a local and global optimization can be run again until a stable and satisfying result is obtained.

### Challenging issues with the dataset

Running conventional machine learning and automated modelling approaches on our dataset is a challenge for several reasons.

1. Its structure and size is strongly unbalanced: 2,741 variables at the genomic scale and 16 at the other scales (i.e. a total of 4 conditions, corresponding to two kinetic points at two different temperatures), with only three replicates for each experimental condition. Finding reliable models is very difficult, as there exist an infinite number of multi-linear models that approximately fit the data.
2. The repeatability of the various measurements is poor (this is rather a common feature for living biological systems).
3. The inter-scale relationships are non-linear [72], a large amount of data is needed to appropriately infer reliable models [73] and avoid over-fitting.

Due to the high variability of the biological data, there is a need to develop a robust model fitting procedure that can easily adapt to different sources of variation. For this purpose, there are several clearly accepted methods for estimating variability or separability, such as the F-test statistic of the one-way ANOVA (Anv) [74], the *p-value* of a Krustal-Wallis test [75], or the Thornton’s separability index [76]. However, although these are well-known statistical methods, their interpretation may be less intuitive.

The F-test statistical method gives more importance to extreme values when the measurements in different conditions are very similar or dissimilar while the *p-value* of the Kruskal-Wallis test corresponds to the probability that the measurements come from the same distribution.

In our case, the objective is not to prove that the results from the different experimental conditions come from similar or different distributions, but rather to provide a qualitative view of the variance existing in the dataset. We thus propose an estimate of the predictive quality of models, as a measure of separability, called *P_V_*, to illustrate the variance existing for each variable *V*, as shown in Eq. 5:

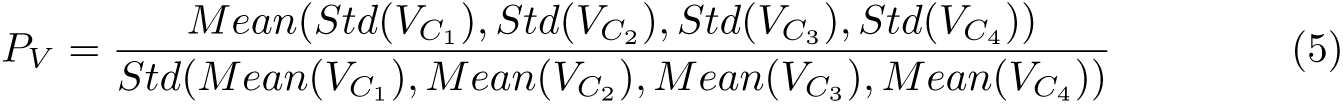

where *V_C__i_* are the 4 replicate measurements of variable *V* for the 4 conditions *C_i_*, and where *Std* and *Mean* are the standard deviation and the mean values respectively. *P_V_* compares the mean of standard deviations of the values for each condition with the standard deviation of the mean value for each condition. Variables for which the distribution of measured values in each condition can be separated, will have a low *P_V_*, near zero, meaning that they can be separated from the rest of the data. Otherwise, *P_V_* tends to infinity. A *P_V_* = 1 means that the mean standard deviation for each condition is as high as the standard deviation of the mean for each condition.

The values of separability *P_V_* for the variables at cellular scales are presented in Table 2. We did not consider the separability of genomic variables because in this application we treated them using clusters and not individually, as described in Step 1 of section Methods. We noted that an average separability value above 1 was obtained for most population-bioprocess scale variables. The only scale where separability is statistically good is the cellular scale where 5 variables among 7 are lower than 0.3. It means that population-bioprocess variables are not separable between the different conditions, in contrast to cell-scale variables. This information is also important in a modelling perspective: It seems that a graph relying on the cell scale variations could be a good solution.

**Table 2.** Separability values for variables from the population-bioprocess to the cellular scale. In the separability value column, a color gradient illustrates the level of data differentiation between the different conditions, ranging from red to indicate a poor separability value (equal to or greater than 1.0) to green to indicate a good separability value (near 0.0).

| Scale | Operating units | Variables | Separability value |
| --- | --- | --- | --- |
| Population-bioprocess | Freezing | dtpH0.7Cong | 1.55 |
|  |  | dtpH0.7speCong | 1.08 |
|  |  | dlog(UFC)Cong | 1.04 |
|  | Freeze-Drying | dtpH0.7Lyo | 0.55 |
|  |  | dtpH0.7speLyo | 0.95 |
|  |  | dlog(UFC)Lyo | 1.57 |
|  | Storage after 3 months | dtpH0.7Sto3 | 2.20 |
|  |  | dtpH0.7speSto3 | 0.71 |
|  |  | dlog(UFC)Sto3 | 1.58 |
| Cellular | Cell membrane | UFA | 0.12 |
|  |  | SFA | 0.20 |
|  |  | CFA | 0.46 |
|  |  | CFA/UFA | 0.10 |
|  |  | CFA/SFA | 0.58 |
|  |  | UFA/SFA | 0.10 |
|  |  | Anisotropy | 0.26 |

## Results

The design of the proposed framework aims at integrating, as much as possible, two types of cognitive knowledge, namely: (i) the deterministic knowledge held in mind by the expert in an explicit form; and (ii) their implicit knowledge that is mobilized when interacting with the computer system. This human-machine collaboration aims at bringing out new hypotheses following the proposal of new or unusual patterns. This way, a biological system can be explored *in silico* by experts with the intention, for example, of designing new experiments aimed at improving the knowledge of the physiological response of LAB to the conditions induced during an industrial bioprocess.

### Overview of the interactive modelling workflow

The exploration workflow is illustrated in Fig 1. As described in Subsection Methods, it is made of two interdependent steps: (i) GIC (for Genomic Interactive Clustering), and (ii) IMSIE (for Interactive Multi-Scale Modelling Exploration).

### Step 1: the GIC method

In this step, clusters are built from the results of a clustering algorithm, the expert then explores the clusters, modify them if necessary, decides if they can be identified as belonging to the same metabolic pathway or as being an operon structure encoding for a biological function, and finally selects the ones that will be used for the model.

In our use-case, from a total of 2,741 variables corresponding to DE genes, using the *L. lactis* TOMSC161 reference genome [56], it was possible to build clusters of genes involved in key metabolic pathways and/or biological functions that may potentially correlate with the physiology of the *L. lactis* TOMSC161 strain under the experimental conditions tested in this study. An overview of the genomic clustering results obtained from 2,741 variables by the GIC method is shown in Fig 2. The interactive interface proposed to the user has four parts:

i. **A dendrogram** representing the hierarchy of nodes with the number of variables that have been clustered (Fig 2A). This step is controlled by the user who defines a minimum size of clusters;
ii. **A schematic representation of the expression profiles** for each gene of the cluster (Fig 2B). When the expert selects a node of the dendrogram, he has access to the content of the corresponding cluster and can quickly evalute is similarity;
iii. **A 2D scatterplot** of the projection of all genes of the dataset using the t-distributed stochastic neighbor embedding (t-SNE) dimension reduction method [77] (this method position each point in a 2D space in such a way that the distances between points correspond to the true distances in the original multidimensional space) (Fig 2C). This is a way to provide another perspective on the clustering results, that may facilitate the analysis and, for instance, can help identify sub-clusters within a node that have not been found by the DB-SCAN hierarchical clustering;
iv. **The list of the names of the variables of the node** (i.e. corresponding to the names of the genes or the locus tag number, as well as the functional product name). The user has the option to remove the variable or variables that seem aberrant with respect to findings from the scientific literature (Fig 2D).

**Fig 2.**
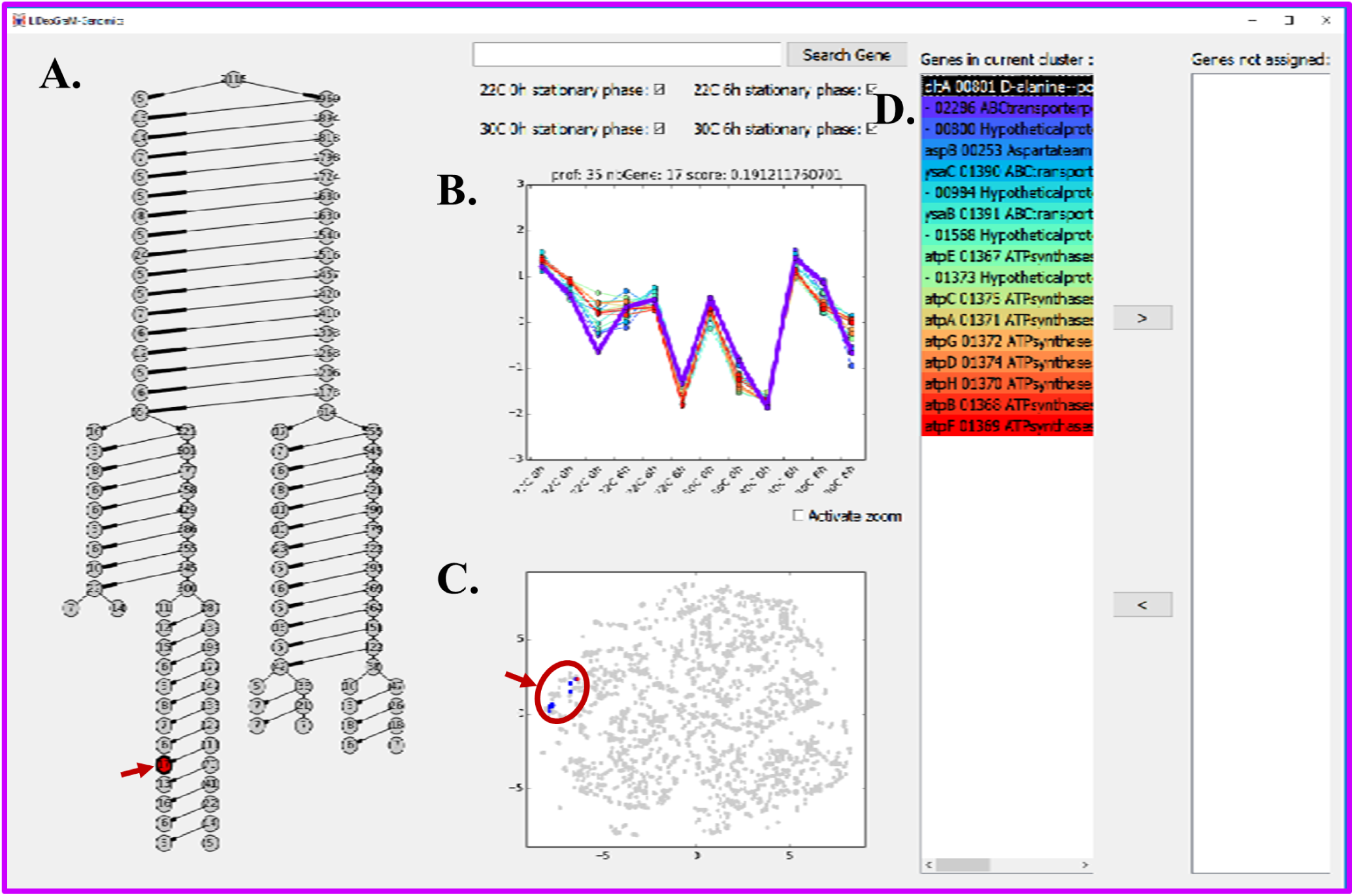
A Screenshot of the workspace during an interactive exploration of the clusters obtained with the GIC method using the 2,741 variables. The graph on the left side (A) represents the hierarchy of clusters. Each node represents a cluster and the number attached to it the number of genes in the cluster. When clicking on a cluster (left, red node) of the dendrogram, the top middle plot (B) shows the standardized expression of the genes in the selected cluster, the bottom middle plot (C) shows the 2D projection of every gene using the t-SNE method, and the associated list of genes is displayed on the right (D). Genes can be assigned or unassigned to a cluster by an expert using the arrows (”*<*” and “*>*”).

If genes are missing in the list for reconstructing the metabolic pathway and/or the associated biological function, the user can issue a query for these genes in the search field using their locus-tag or name. For each returned gene, it is added to the list and to the lowest hierarchical cluster in which it appears. Conversely, if a gene erroneously appears in the list, it can be removed from the list and from the cluster.

### Step 2: the IMSIE method

A graph structure is first built to setup some constraints (explicit expert knowledge) on the potential links between scales (step 2.1). Then a multi-linear regression algorithm (see section Methods) proposes a set of possible local models according to the previous constraints.

An interactive exploration of the proposed models is triggered (step 2.2) to let the expert select and revise some appropriate set of local models (implicit expert knowledge). We have chosen to strengthen the expression of the human knowledge at the local level, variable by variable, because it has been shown that knowledge is often more easily mobilized at this level. A reason for that is that laboratory experiments are often guided by local laws. The challenge was also to drive the experts to express their non-explicit knowledge through this concrete exploration and is inspired from the methodology of elicitation proposed by [78], already applied and validated in previous studies [79].

Once the expert is satisfied with the local hypothesis, a stochastic optimization algorithm (see section Methods) then builds a coherent global model by retaining only one local model per variable so that the available data are best predicted by the chained local models (step 2.3). All local and global models can be reviewed and the corresponding machine learning and optimization procedures can be restarted as necessary until a global model that satisfies the expert is obtained. More details are presented below in a step by step fashion.

### Step 2.1: Interactive setup of graph constraints

The interface proposed to the user and presented in Fig 3 is interactive. The user is free to create the connection links, to add or remove them between the different categories of multi-scale variables (i.e. population-bioprocess, cellular and genomic, (Table 1)), which are then considered in the prediction. Each of the links, as well as all the names of selected genomic clusters, are listed in a dedicated history on the left and right of the local window, respectively (Fig 3), to keep the user informed of the choices made. The expert specifies the permitted, mandatory or prohibited links between variables and groups of variables (i.e. clusters from step 1) at different scales (Fig 3). The possibility to manipulate variables grouped into coherent clusters and individual variables helps reduce complexity [79, 80], which is helpful in assisting reasoning across multi-scales. In this interface, a graph represents the structure of the multi-scale model in a stratified way, where each node is associated to a group of variables, and each connection corresponds to the possibility of expressing variables associated to the child node as a function of the variables associated to the parent nodes.

**Fig 3.**
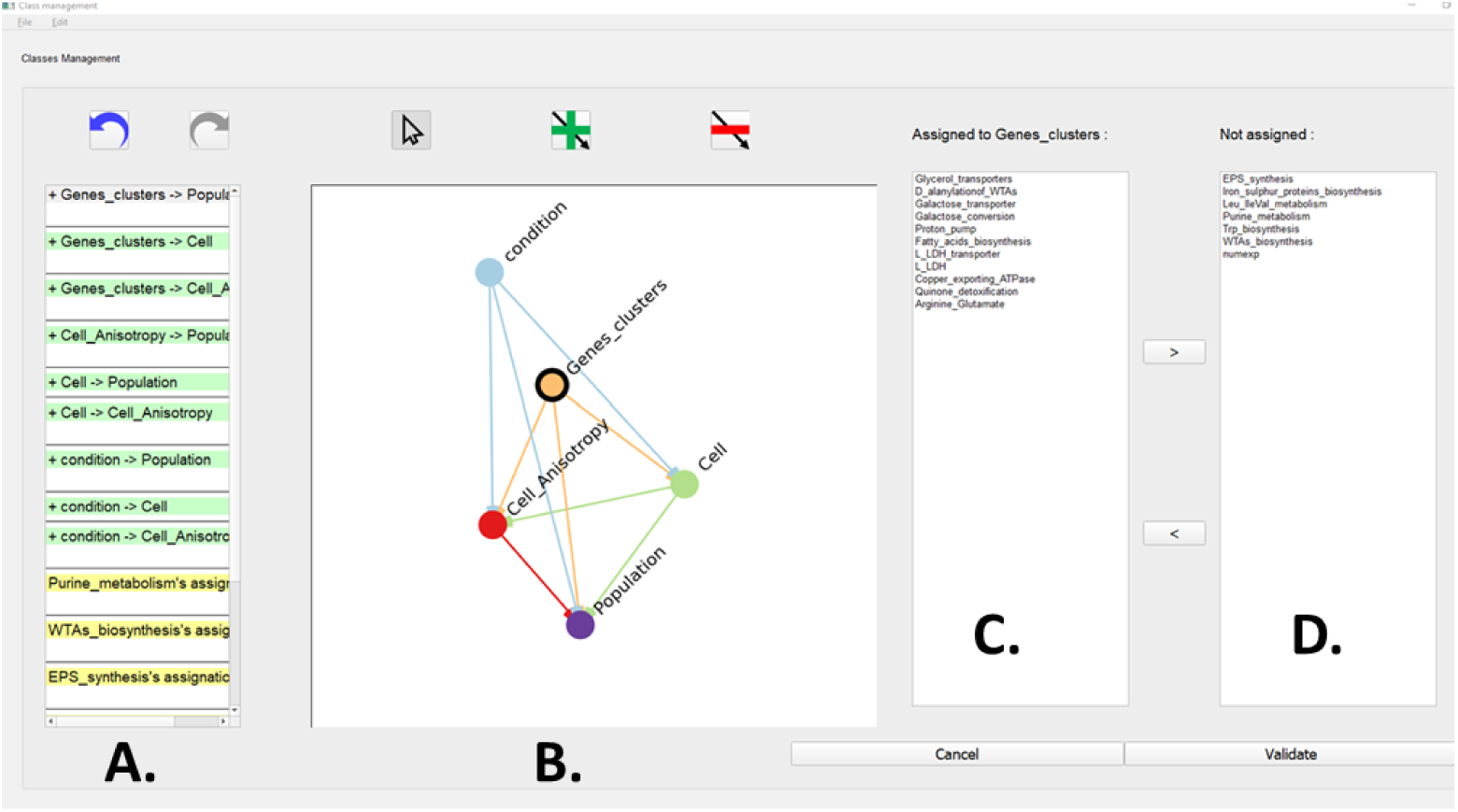
A screenshot of IMSIE’s interactive manager for building constraints. (A) **A history of currently selected constraints.** (B) **Graph representation of the variables (nodes) and contraints (edges). The blue node called “condition” represents the experimental conditions. The red and green nodes called “CellAniso” (for the anisotropy component) and “Cell” (for the membrane fatty acids component) respectively, represent the cellular scale. The purple node represents the population scale and the orange node with a black outline called “Genes” represents the genomic scale. An arrow corresponds to a constraint set by the user, added and removed with the ‘+’ and ‘-’ buttons respectively (top icons). The color and direction of the arrows indicate the direction of the constraint (the output node depends on the input nodes). All the arrows are listed in the history. A node selected by the user (mouse pointer) is displayed with a black outline.** (C) **The set of gene clusters associated with the selected variable.** (D) **The set of gene clusters that are not yet assigned with a variable.**

Using this interface it is thus possible to review the grouping of variables, to delete or create new groups. Variables can also be removed in order to ignore them in the following steps. In our use-case, the constraints imposed between the different scales were established in a hierarchical framework that links the smallest scale (i.e. genomic) to the largest scale (i.e. population/bioprocess). For example, it is possible to: (i) connect the category of variables representing the genomic scale (referred to as “Genes clusters”) to the categories of variables representing the other scales; (ii) connect the categories of variables representing the cellular scale (referred to as “Cell” and “CellAniso”) to the categories of variables representing the population scales; or even (iii) prohibit links, like for example Fig 3 where the node “condition” is not linked to the node “genes”, inducing that node “genes” as another input of this model.

In addition, it should be noted that in these proposed constraints, the measurement of the cell’s Anisotropy (referred to as “CellAniso”) and the set of different fatty acid components of the cell membrane (referred to as “Cell”) both representing the cellular scale, were separated by experts into two distinct categories for biological signification purposes [51].

This first exploration step allows experts to express their knowledge in an explicit way (typically in less than 30 minutes), to establish an initial model.

### Step 2.2: Interactive shaping of local models

After computing the local models using a multilinear regression (see section Methods), an interface is presented to the expert for exploration and interaction (Fig 4A). The interface is designed to enable a multi-faceted exploration using coordinated multiple views [81], and highlights the parts of the multi-scale model that needs expert attention as previously described in [68]. As such, the equations are accessible in three different forms through multiple coordinate views following:

i. a graphical view showing the average fit of local models obtained by symbolic regression (Fig 4A); Each node of the graph **(a)** corresponds to a variable or a group of variables, as selected by the user in step 2.1 (Fig 3), each arrow indicates the existence of at least one equation between its extremities, and the colours give a global evaluation of the average quality of local models. According to the choice of the user, different types of information are displayed when clicking on a node or a link;
ii. a table with five levels of complexity, displaying the proposed mathematical equations for a selected node (Fig 4A)**(b)**; For each equation, three measures are displayed: complexity, fitness score (0 corresponds to a perfect prediction) and “random” fitness (a high value means that the considered equation is not better than a random prediction). These values are highlighted in colours ranging from green (good) to red (poor);
iii. an overview of the scatterplot on a linear regression curve, informing us about the quality of correlations between the predicted and measured data for the selected local model (Fig 4A)**(c)**.

**Fig 4.**
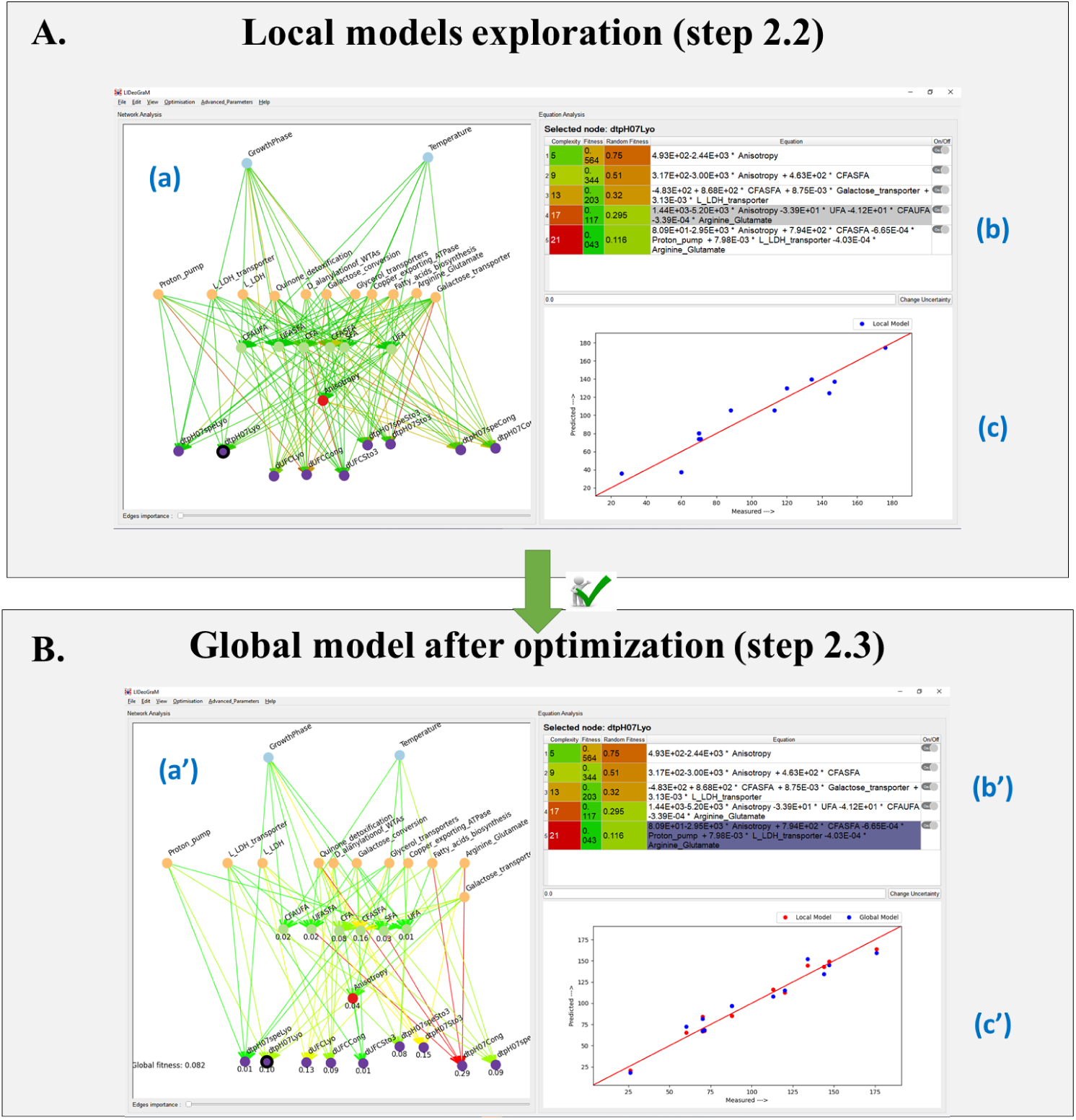
Screenshots of Step 2 interface widgets. **A:** construction of local models, **B:** design of the global model. **(a)** and **(a’)**: graph representations of local and global models. Each node corresponds to a variable or a group of variables, they are coloured according to scales: bioprocess (i.e., blue nodes), genomic (i.e., orange nodes), cellular (i.e. green and red nodes) and population (i.e. purple) scales. The color of the arrows corresponds to their predictability, ranging from green (good predictability) to red (poor predictability). The selected node is outlined in black. **(b)** and **(b’)** display the equations associated with the selected node. The predictive quality of each equation is indicated by a color gradient from red (close to 1, poor prediction) to green (close to 0, good prediction). The equation selected by the expert is indicated in gray (b) for the local models. In purple (b’) is underlined the equation selected for the global model after optimization. **(c)** and **(c’)** display the scatterplots of predicted versus measured data (the red line corresponds to a theoretical perfect prediction). In (c’), local (red dots) and global (blue dots) models are displayed together to allow for comparison.

All steps can be repeated as desired, including the calculation of new sets of equations. We observed that over-constraining step 2.1 appears to have detrimental results. A balance must be progressively found, allowing the user to test, accept, or reject different hypotheses.

### Step 2.3 Interactive design of the global model

A global model is then computed and displayed in a similar way (Fig 4B). This global model uses only one equation per variable (highlighted in purple), so that the output variables are calculated from the input variables using a cascade of local models.

The user can modify the global model, replace a selected equation by another and test it against measured data. It is possible to return to the local model’s interface component (widget), re-run new calculations and again generate a new global model.

#### Remark

The predictive modelling tool we propose, made up of 3 sub-steps, has multiple advantages compared to existing approaches [3]:

- *A priori* explicit knowledge is balanced with *a posteriori* implicit knowledge, through a back-and-forth approach between local and global perspectives on the system;
- Step 2.2 is a convenient tool for introducing a diversity of models, on which the implicit knowledge of experts can be mobilized;
- The comparative visualization of the results of local and global models tends to limit the propagation of errors (due to noisy and/or erroneous data, for example);
- Experts can remove invalid edges or revise nonsensical equations, which also reduces the propagation of errors.

### Proof of concept: analysis of a multi-scale engineering application with *Lactococcus lactis* TOMSC161, a LAB of industrial interest

Given the multi-disciplinary nature of the proof of concept use-case described below, multiple types of expertise were needed to explore and validate the clustering results and the different models generated by **Biosys-LiDeOGraM**. We thus collaborated with two experts from functional genomics and two experts from process engineering. These experts were senior researchers with, on average, over 20 years experience in their respective domains. They participated in this project, from data collection, integration and structuring, to exploration (clustering, and multi-scale modelling), analysis and interpretation of the results.

The use-case exploration and analysis took place between 2020 and 2021 (8 months period), where regular meetings were held often lasting 30—45 minutes each. **Biosys-LiDeOGraM** was displayed using a laptop (13” screen), a standard desktop monitor (19”) or a large tactile display (65” UHD-4K). In total there were around 25 exploration sessions, held either in person or remotely (due to the COVID pandemic). During the first session, a brief demo of the tool was provided to explain its main functionalities. In each session, a single domain expert conducted most of the interaction with the tool, while the other domain experts discussed without interacting.

The observations and findings we report in the use-case below are based on notes taken during these meetings, as well as various artifacts collected during the collaboration, such as screenshots of the tool, mind maps of conceptual discussions, and system log files. Additionally, the functional genomics expert subsequently carried out different types of analyses and interpretations of the experiment results, particularly for Figures 7 and 8, to verify if the proposed correlations between the scales made biological sense.

### Interactive model design from data

Fig 5 shows a summary of the modelling process: after about ten iterations, 17 of the 25 genomics clusters generated from 2,741 DE genes variables by the GIC method were used in order to: (i) save computing time (approximately a gain of 30 minutes per variable) (ii) preserve the cross-scale complexity; (iii) limit the random proposals per variable; and (iv) maintain consistency of the selected metabolic pathways and/or biological functions with the other scales when it is known (developed below in the IMSIE exploration step). The 7 variables at the cellular level and 9 variables at the population-bioprocess level (described in Table 1) were kept in the final model.

**Fig 5.**
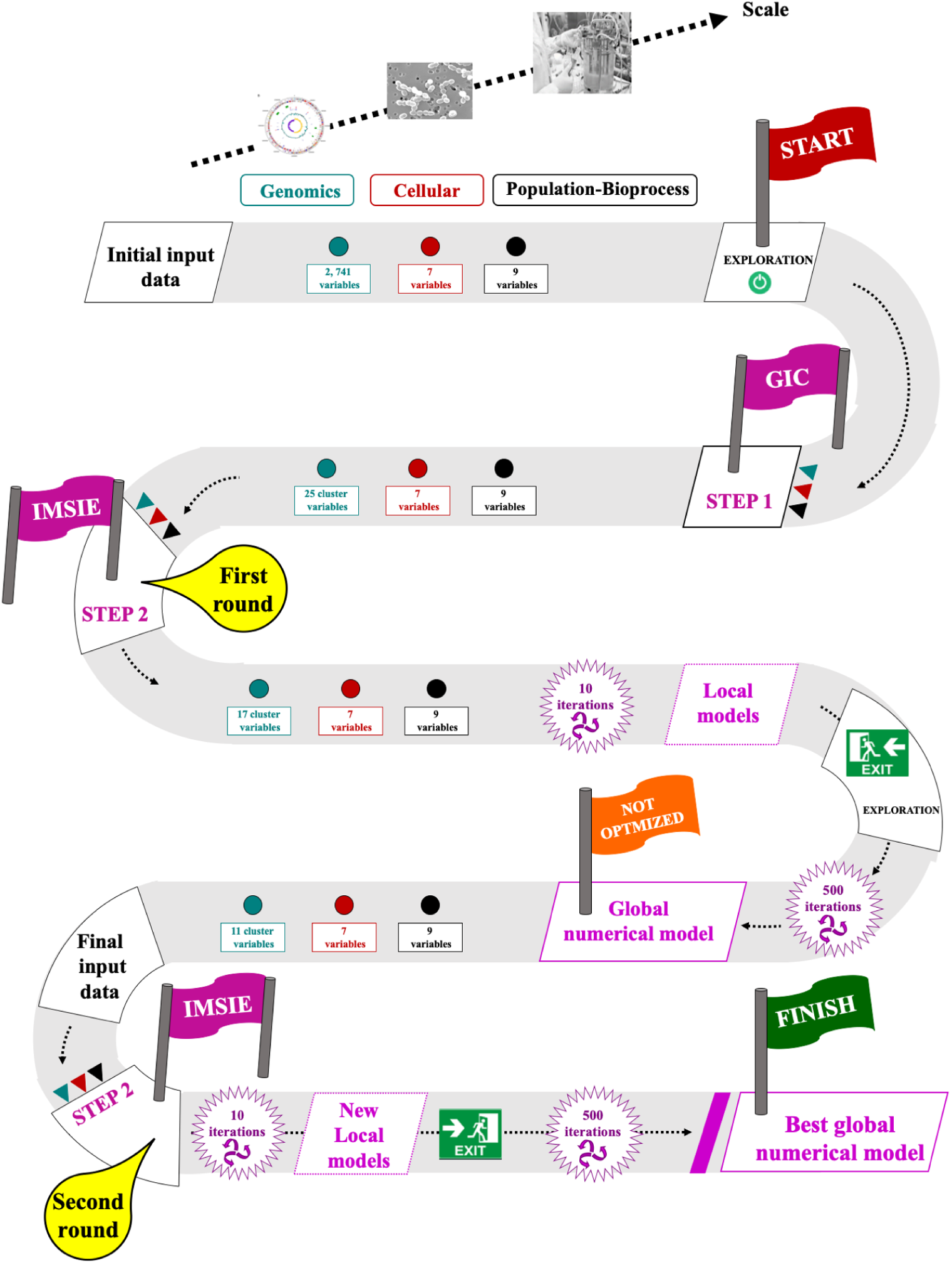
The variable and space reduction path for local and global models using Biosys-LiDeOGraM. The cyan, red, purple and black dots illustrate the genomic, cellular, population and population-bioprocess datasets, respectively. The number of variables of each dataset at different scales is given. The same color code is used for triangles, which represent input variables for step 1 (GIC algorithm) and step 2 (IMSIE algorithm).

### Genomic interactive clustering (GIC) exploration

The objective of this exploration is to gain an in-depth understanding of how the genomic scale could influence the biological properties and behavior of LAB at higher scales, which would help us to preserve their functional properties by choosing the best control parameters for the fermentation, freezing and freeze-drying processes. For this purpose, we choose to work with transcriptomic data as it largely reflects the overall response of a microorganism under given environmental stress conditions.

Like all clustering tools, our approach is not perfect. Uncertainties and/or errors often appear when assigning variables to clusters. To reduce interpretation errors, an initial clustering stage is complemented by an additional molecular genetics analysis step. This method was performed on 35 nodes having between 2 and 18 variables. The restrictive choice was made because nodes with more than 20 variables, as well as those with less than two variables, are not discriminant enough to be correlated with metabolic pathways or biological functions. Subsequently, several analysis criteria were applied to these nodes in order to eliminate the genomic clusters that are independent of our study, and therefore would lead to erroneous interpretations if integrated into the multi-scale model. Thus we discarded the genomic clusters corresponding to: (i) clusters of prophage regions; (ii) clusters of ribosomal translational genes; (iii) clusters of unconnected genes; (iv) incomplete clusters (i.e. corresponding to more than 50% of the missing genes for the associated biological function); and/or (v) genes expression profiles that are different for a single cluster.

Ultimately, only 25 genomic clusters generated by the GIC method were validated and selected to contribute to the multi-scale model. Their genetic characteristics and their biological functions are presented in Table 3.

**Table 3.**
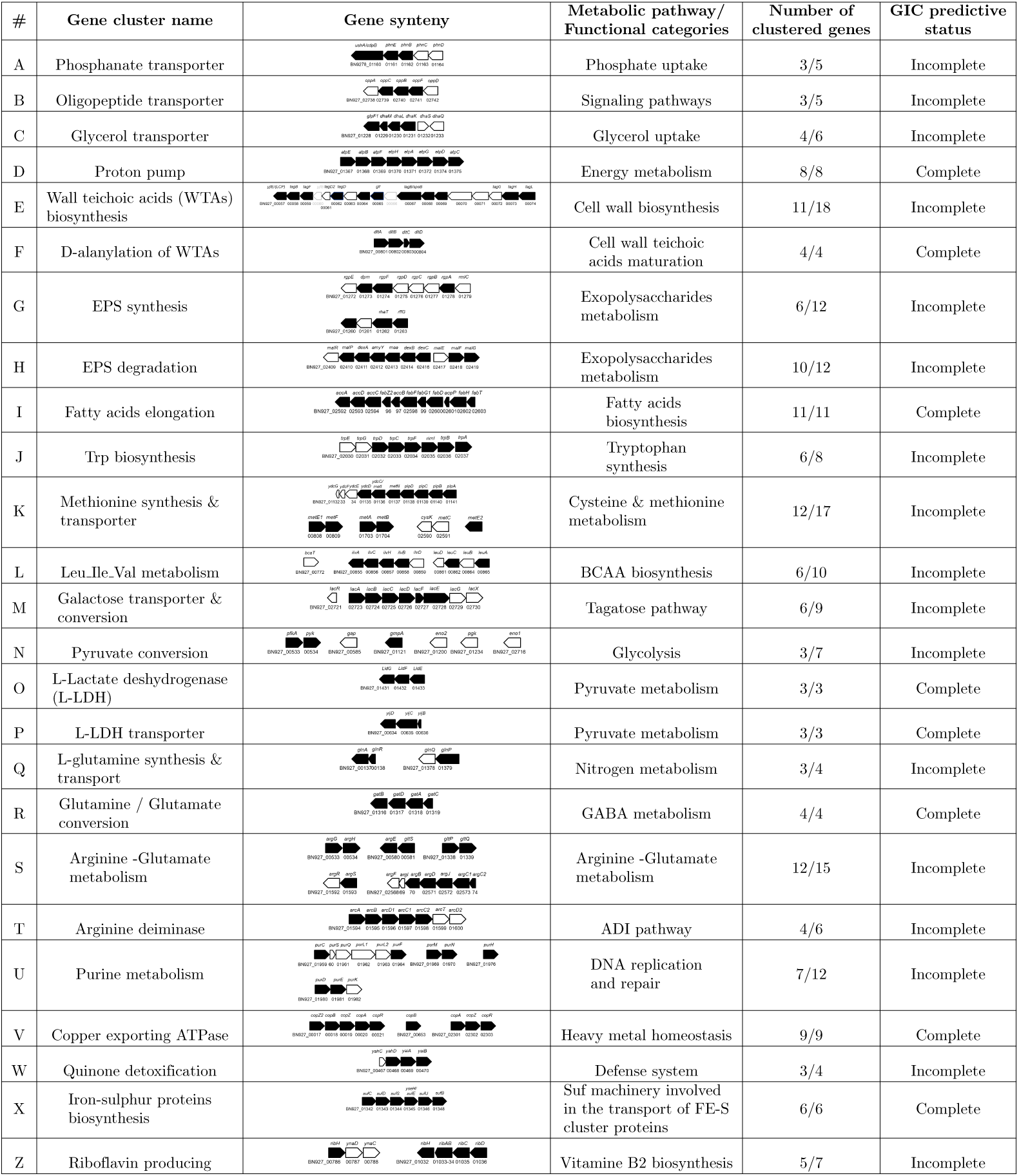
The list of the 25 genomic clusters and their characteristics, generated by the GIC method and validated by the expert. In the genetic environment, the black and white pentagons represent respectively detected and undetected genes associated with the metabolic pathway and/or biological function. The number of genes detected by the GIC predictive method out of the original number of genes in the genetic cluster is indicated for each associated metabolic pathway and/or biological function.

| # | Gene cluster name | Gene synteny | Metabolic pathway/<br>Functional categories | Number of<br>clustered genes | GIC predictive<br>status |
| --- | --- | --- | --- | --- | --- |
| A | Phosphatase transporter |  | Phosphate uptake | 3/5 | Incomplete |
| B | Oligopeptide transporter |  | Signaling pathways | 3/5 | Incomplete |
| C | Glycerol transporter |  | Glycerol uptake | 4/6 | Incomplete |
| D | Proton pump |  | Energy metabolism | 8/8 | Complete |
| E | Wall teichoic acids (WTAs) biosynthesis |  | Cell wall biosynthesis | 11/18 | Incomplete |
| F | D-alanylation of WTAs |  | Cell wall teichoic acids maturation | 4/4 | Complete |
| G | EPS synthesis |  | Exopolysaccharides metabolism | 6/12 | Incomplete |
| H | EPS degradation |  | Exopolysaccharides metabolism | 10/12 | Incomplete |
| I | Fatty acids elongation |  | Fatty acids biosynthesis | 11/11 | Complete |
| J | Trp biosynthesis |  | Tryptophan synthesis | 6/8 | Incomplete |
| K | Methionine synthesis & transporter |  | Cysteine & methionine metabolism | 12/17 | Incomplete |
| L | Leu.Ile.Val metabolism |  | BCAA biosynthesis | 6/10 | Incomplete |
| M | Galactose transporter & conversion |  | Tagatose pathway | 6/9 | Incomplete |
| N | Pyruvate conversion |  | Glycolysis | 3/7 | Incomplete |
| O | L-Lactate deshydrogenase (L-LDH) |  | Pyruvate metabolism | 3/3 | Complete |
| P | L-LDH transporter |  | Pyruvate metabolism | 3/3 | Complete |
| Q | L-glutamine synthesis & transport |  | Nitrogen metabolism | 3/4 | Incomplete |
| R | Glutamine / Glutamate conversion |  | GABA metabolism | 4/4 | Complete |
| S | Arginine -Glutamate metabolism |  | Arginine -Glutamate metabolism | 12/15 | Incomplete |
| T | Arginine deiminase |  | ADI pathway | 4/6 | Incomplete |
| U | Purine metabolism |  | DNA replication and repair | 7/12 | Incomplete |
| V | Copper exporting ATPase |  | Heavy metal homeostasis | 9/9 | Complete |
| W | Quinone detoxification |  | Defense system | 3/4 | Incomplete |
| X | Iron-sulphur proteins biosynthesis |  | Suf machinery involved in the transport of FE-S cluster proteins | 6/6 | Complete |
| Z | Riboflavin producing |  | Vitamine B2 biosynthesis | 5/7 | Incomplete |

Even if the result of GIC remains perfectible (only 32% of the genomic clusters are complete), the current clustering is however acceptable since we found that the distribution of each variable within a genomic cluster was correctly assigned, and as we will see below, it has allowed us to reconstruct the associated metabolic pathway and/or biological function. In addition, all selected clusters were considered important to explain the physiological behavior of *L. lactis* TOMSC161 in the fermentation, freezing and freeze-drying processes.

### Interactive multi-scale modelling (IMSIE) exploration

This process starts with a preliminary model calculated without expert interaction: a first complete graph of constraints was tested, linking all the variables except the temperature and growth-phase variables since they are considered model input and with the 17 genomic cluster variables out of 25, considered as essential by the expert for this exploration. The result (Fig 6 A.) was predominantly composed of red arrows, indicating poor prediction for the majority of the nodes (27/33 nodes). The global fitness score was 0.777 (close to 1, thus a rather poor result). At the population-bioprocess scale in particular, we obtained a poor prediction of nodes with an average fitness score of 0.841 +/- 0.185. This suggests that the chosen constraints linking the experimental conditions: “Temperature”and “Growth-Phase” with one of the 17 selected genomic clusters do not permit correlation with the fermentation, freezing/freeze-drying and storage processes. Similarly, at the genomic scale, the performance was poor for each of the nodes (fitness scores close to 1.00 except for the cluster “Glycerol transporters”), meaning that these results were obtained randomly. Conversely, we noticed a relatively good predicted value for 4 nodes out of the 6 cell membrane variables (“SFA”, “CFA/UFA”, “UFA/SFA” and “UFA”) at the cellular scale with a fitness score close to 0, ranging from 0.03 to 0.10. For the 2 cellular scale variables (“CFA/SFA” and “CFA”), the use of the genomic cluster component strongly decreases the quality of node prediction, with a score close to 1. Given these prediction results and the computed separability values (see section Methods) for the metabolic pathways and/or biological functions selected by the experts (Table 4), we concluded that the integration of the 17 selected genomic clusters is probably too restrictive considering the sparsity of the available data.

**Fig 6.**
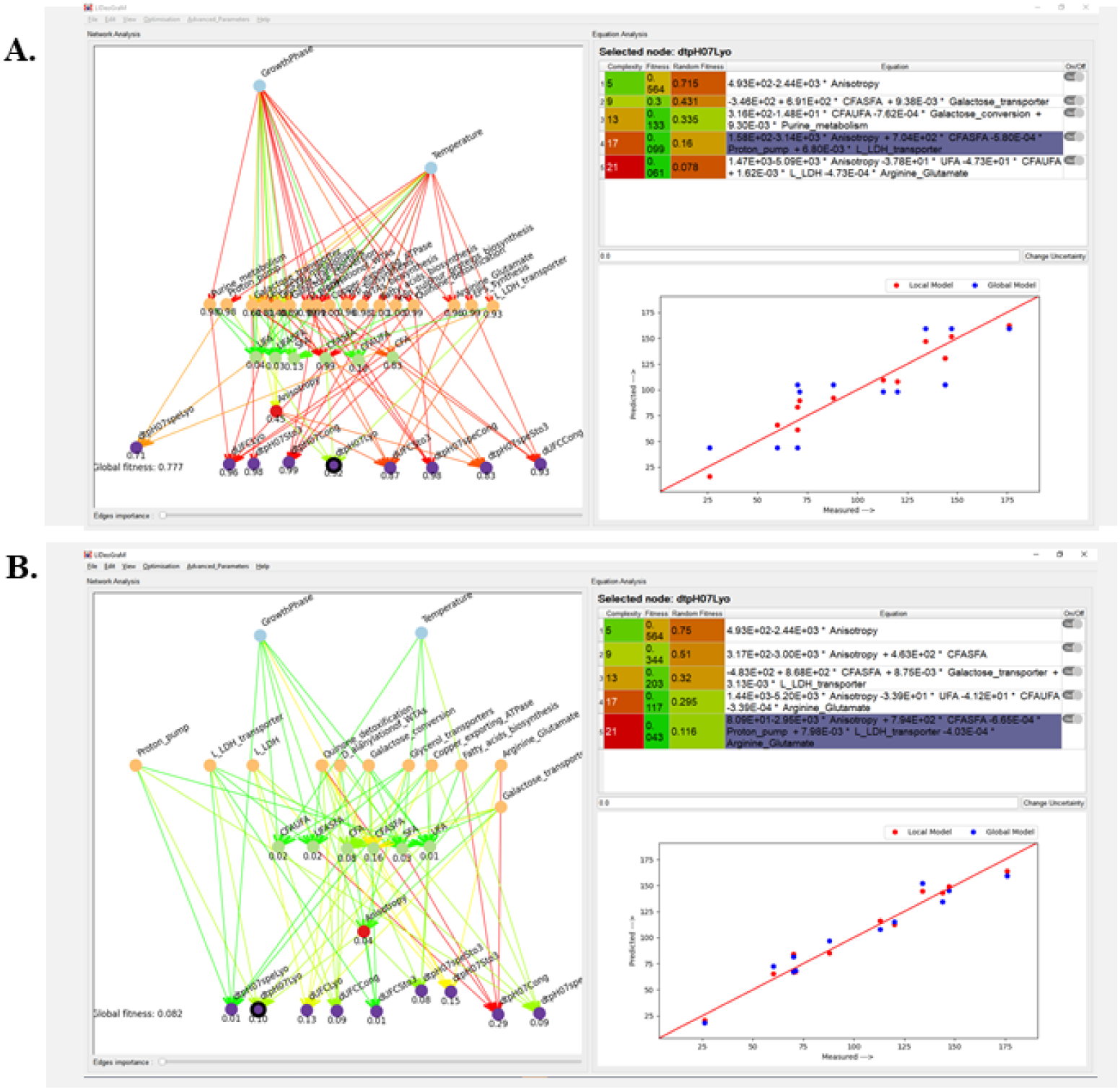
The result of the global model without (A.) human-machine interactions and after several rounds of human-machine interactions (B.).

**Table 4.** Computed separability values for the genomic scale variables as selected by experts. Values in the second column from the left correspond to the number of genomic clusters selected for the first exploration (where a total of 17 genomic clusters were selected). The numbers with grey background correspond to the genomic clusters that were removed for the second exploration (a total of 11 genomic clusters are kept). In the separability value column, a color gradient illustrates the level of data differentiation between the various conditions, ranging from red to indicate a poor separability value (equal to or greater than 1.0) to green to indicate a good separability value (near 0.0).

| Scale | # | Selected variables<br>(Gene cluster name) | Separability value |
| --- | --- | --- | --- |
| Genomic | 1 | Leu-Ile-Val metabolism | 0.42 |
|  | 2 | Glycerol transporter | 0.30 |
|  | 3 | D-Alanylation of WTAs | 1.87 |
|  | 4 | WTAs biosynthesis | 1.10 |
|  | 5 | EPS biosynthesis/degradation | 1.64 |
|  | 6 | Galactose transporter | 0.24 |
|  | 7 | Galactose conversion | 1.13 |
|  | 8 | Fatty acids elongation | 3.69 |
|  | 9 | Proton pump | 0.83 |
|  | 10 | L-lactate deshydrogenase (L-LDH) | 1.06 |
|  | 11 | L-LDH/substrate transporter | 0.88 |
|  | 12 | Copper exporting ATPase | 1.07 |
|  | 13 | Quinone detoxification | 1.42 |
|  | 14 | Trp biosynthesis | 1.47 |
|  | 15 | Arginine - Glutamate | 1.07 |
|  | 16 | Iron-sulphur proteins biosynthesis | 1.21 |
|  | 17 | Purine metabolism | 1.98 |

Consequently, we chose to improve node prediction by: (i) forbidding the links between “temperature” and “age/growth phase”, and the selected genomic clusters; (ii) working only with 11 of the 17 genomic clusters (Table 4) that seem to be more significant for the domain expert; and finally (iii) enforcing the links between “anisotropy” and these 11 clusters. A new exploration with IMSIE then yields the results shown in Fig. 6B. The resulting network is now predominantly green with a global fitness score of 0.082 and a perfect linear correlation between measured and predicted data for the majority of local and global models. In particular, we obtained very good fitness scores for both the “cell membrane” and “anisotropy” variables (ranging from 0.01 to 0.16) with the “temperature *∗* age/growth phase” and any of the 11 selected genomic clusters at the cellular scale, indicating a strong correlation as expected by the experts. Similarly, at the population-bioprocess scales, we noticed a clear improvement over the first global model that was proposed without constraints (Fig. 6A), for the prediction of the 9 associated nodes with an average fitness score of 0.105 + */ −* 0.083, reflecting a fairly good integration of the different multi-scale variables by the machine-learning algorithm, within the imposed constraints.

### Summary

This use-case shows that **Biosys-LiDeOGraM** was able to: (i) engage domain experts in the clustering and model exploration steps which helped improve the final model; (ii) integrate biological data from various experiments at different scales even in the presence of limited data; and (iii) establish valid correlations between the different multi-scale variables (as shown in the next section).

### The proposed model of physiological response of the LAB

#### Biological analysis of the proposed equations

For validating the model built with **Biosys-LiDeOGraM**, we analyzed the equations of 11 key nodes as identified by the domain expert. These equations, highlighted in blue/purple color in Fig 7 were selected by the final optimization procedure and thus are part of the final global model (100 iterations of the evolutionary algorithm was considered sufficient when compared to an initial run test with 500 iterations).

**Fig 7.**
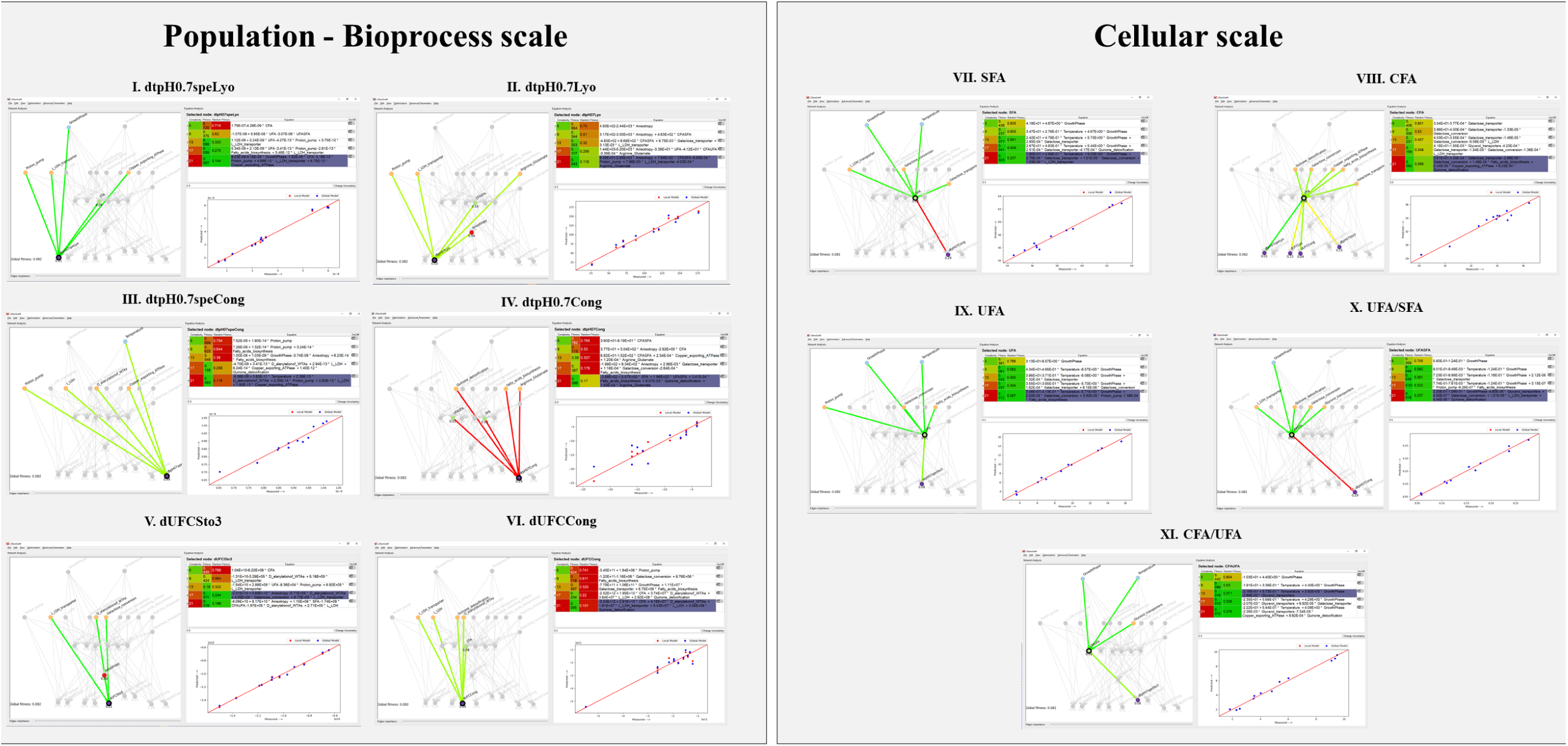
Validation of the 11 selected nodes at different multi-scale levels. We show the predictions of local models for the nodes “dtpH0.7speLyo”, “dtpH0.7Lyo”, “dtpH0.7speCong”, “dtpH0.7Cong” (**I** to **IV**) of the population-bioprocess scale, “dUFCSto3”, “dUCFCong” (**V** and **VI**) of the population scale, and “SFA”, “CFA”, “UFA”, “UFA/SFA”, “CFA/UFA” (**VII** to **XI**) of the cellular scale. Good quality predictive models are coloured in green, while poor quality ones are in red. The five proposed equations are highlighted in purple, and the corresponding scatterplots of the predicted versus measured data for local (red dots) and global (blue dots) models are displayed.

At the population-bioprocess scale, the local models for 4 selected nodes, “dtpH0.7speLyo”, “dtpH0.7Lyo”, “dtpH0.7speCong” and “dtpH0.7Cong” (**I** to **IV**) in Fig 7, were considered to explain the observed variable at the population scale. For the “dtpH0.7speLyo” node, which represents the overall biological activities of starters LAB in the freeze-drying process, the local model reaches a good prediction score (fitness value of 0.001), and indicates a dependence on the “age/growth phase” component, the 3 gene clusters: “proton pump”, “L-LDH substrate transporter” and “copper exporting ATPase” for the genomic component, and the membrane fatty acid “CFA” of the cellular component. From a biological point of view, these dependencies are consistent with the knowledge of LAB physiology. It is clear that the components corresponding to the “age/growth phase”, the “proton pump” and “L-LDH substrate transporter” gene clusters are correlated with the overall biological activities of starters at the end of the fermentation process. Indeed, the production of lactic acid/lactate by the bacterial cells is exported to the culture medium by the L-LDH substrate transporter resulting in a decrease in the pH. This acidifying activity induces physiological changes in bacterial cells. Consequently, it is necessary for cells to maintain the pH homeostasis to preseve cell viability, in particular through the action of the proton pump. Furthermore, the “copper exporting ATPase” is strongly correlated with changes in the “CFA” ratio of the cell membrane components. This is not surprising, since the maintenance of heavy metal homeostasis is essential for cell viability, particularly under stressful conditions to limit the potential deleterious phenomenon of membrane lipid peroxidation [50].

Except for the selected “dtpH0.7Cong” node, the variables incorporated into the predictive local models for the “dtpH0.7Lyo” and “dtpH0.7speCong” nodes share similarities with the “dtpH0.7speLyo” node, including the genomic component correlated with acidifying activity, and additional variables that were correlated with cell integrity during the freeze-drying and/or freezing process. For the “dtpH0.7Cong” node, we noticed that the prediction of the local model is less accurate (local fitness score of 0.29 at the highest level of complexity, Fig 7), and remains optimized with more informative data or using non linear equations. However, the variables proposed in the equation that predict this variable are far from being outliers. Indeed, at the genomic scale, “the Fatty acid biosynthesis” gene cluster is directly involved in the lipid composition of the cell membrane, and therefore it is not surprising that it is coupled with the “SFA” and “UFA/SFA” variables at the cellular level. Furthermore, the addition of the “Arginine Glutamate” gene cluster and the “Quinone detoxification” gene cluster in the genomic component may also explain the physiological state of LAB under our experimental conditions. In the related literature, it is clearly recognized that arginine and glutamate which provide an additional source of energy, as well as carbon and nitrogen, which are necessary for LAB growth, all play a key role in maintaining pH homeostasis, thereby improving cell tolerance to acidic conditions [24, 82]. Probably, bacterial cells face toxicity from copper-induced quinones, but this is counteracted by a detoxification system, reducing cell viability loss post-freezing [83]. Although the local model appears to capture available knowledge, it falls short in optimizing the global model, likely due to insufficient data for predicting other levels.

For “dUFCSto3” (**V** in Fig 7), reflecting the loss of cell viability after three months of storage, the good predictions of the local model (fitness value of 0.01) indicate that this variable depends on the 2 gene clusters: “L-LDH” and “D-alanylation of WTAs” for the genomic component, and on the variables “anisotropy”, “SFA” and “CFA/UFA” for the cellular component. One hypothesis to explain this would be that the bacterial cells probably induce changes in their cell envelopes as a result of their acidifying activity via the action of L-LDH enzyme that converts pyruvate to lactate during the fermentation process. This is evidenced by the “D-alanylation of WTAs” gene cluster’s involvement in cell-wall maturation [84] and modifications in cell membrane fatty acid composition, specifically “SFA” and “CFA/UFA”. The changes in the fatty acid composition and ratios lead to a decreased membrane fluidity, as measured by “anisotropy”, improving tolerance during storage. Similarly for the “UFCCong” node (**VI** in Fig 7), we found common variables at the genomic and cellular levels which were previously described at the population-bioprocess level. These variables are closely correlated with cell viability, and be consistent with the freezing process.

At the cellular scale, the local models for 5 selected nodes: “SFA”, “CFA”, “UFA”, “UFA/SFA” and “CFA/UFA” (**VII** to **XI** in Fig 7) were considered to highlight existing relationships between physico-chemical conditions, gene clusters and/or processes related to cell membrane characteristics. We noticed that regardless of the nodes considered, with the exception of the “CFA” node, the fatty acid composition is directly related to “temperature” and/or “growth phase” components. In fact, it is clearly recognized that during fermentation, bacterial cells are able to modulate the composition of their lipid membrane according to the temperature, pH variation, composition of the culture medium in order to preserve vital functions such as: assimilation and conversion of carbohydrates, transport and/or diffusion of solutes, and other metabolic pathways involved in the physiological response of LAB [25]. It is therefore not surprising to observe that 4 out of 5 of these local predictions include the genomic component’s gene clusters: “galactose conversion” and/or “galactose transporter”, involved in *L. lactis* sugar fermentation and contributing to acidifying activities [85]. In addition, the local predictive models for the variables “SFA”, “UFA” and “UFA/SFA” (**VII, IX** and **X** in Fig 7), indicate an additional genomic component that corresponds to the “L-LDH transporter”, “proton pump” and/or “Fatty acid biosynthesis” gene clusters. These close relationships are likely due to the pH change caused by the acidifying activities of LAB, affecting the ratio of saturated and unsaturated fatty acids through the fatty acid biosynthesis pathway. This leads to changes in cell membrane properties [86], causing a loss of cell viability in the freezing, freeze-drying, and storage processes.

Finally, we found that “copper-exporting ATPase” and/or “quinone detoxification” gene clusters play an important role in the locals model of “CFA” and “UFA/SFA” nodes (**VIII** and **XI** in Fig 7). As previously mentioned at population-bioprocess level, bacterial cells appear to be stressed by the toxicity of copper and quinone, which again can be lethal to cells. Thus to ensure their survival, cells maintain copper homeostasis and trigger detoxification systems that limit the degradation of membrane unsaturated fatty acids [83].

To conclude, in this use-case, correlations were established to explain the tolerance or loss of cell viability of LAB during freezing, freeze-drying and storage processes. This was made possible thanks to a collection of nested local models involving multi-scale variables, built using machine learning.

#### A coherent computational model for the physiological responses of *L. lactis* to stress

LAB starters are major players in the manufacture of fermented foods. To be useful, these bacteria must withstand stress conditions that are sometimes drastic especially during fermentation and stabilization processes (freezing, freeze-drying). These conditions can harm the cell envelope, affecting its cellular physiology and enzymatic activities, as well as decrease its stability, resulting in loss of viability [50].

However, microorganisms show remarkable adaptive capacities to certain physicochemical stresses encountered in their natural environment. The triggering of complex molecular mechanisms generates a physiological response of the microorganism that leads to a state of tolerance, and consequently to its survival under normally lethal conditions [24, 50]. In this context, tolerance to abiotic stresses in LAB appears to be a key factor for the control of the quality of the final product.

Using the equations proposed by **Biosys-LiDeOGraM**, at different population-bioprocess and cellular scales (Fig 7), a global numerical model of the physiological response from *L. lactis* could be established and is summarized in Fig 8. A total of 9 common gene clusters involved in key metabolic pathways and biological functions were found in the different equations, enabling tracing of the physiological response of *L. lactis* from the molecular to the population-bioprocess levels. During fermentation, LAB metabolize sugars, mainly lactose via the glycolysis pathway and also through the tagatose pathway, which involves the transport and conversion of galactose to pyruvate. This pyruvate is then converted to L-lactate by L-lactate dehydrogenase (*L-LDH*), supplying energy for optimal growth and producing microbial biomass and acidifying activities (Fig 8).

**Fig 8.**
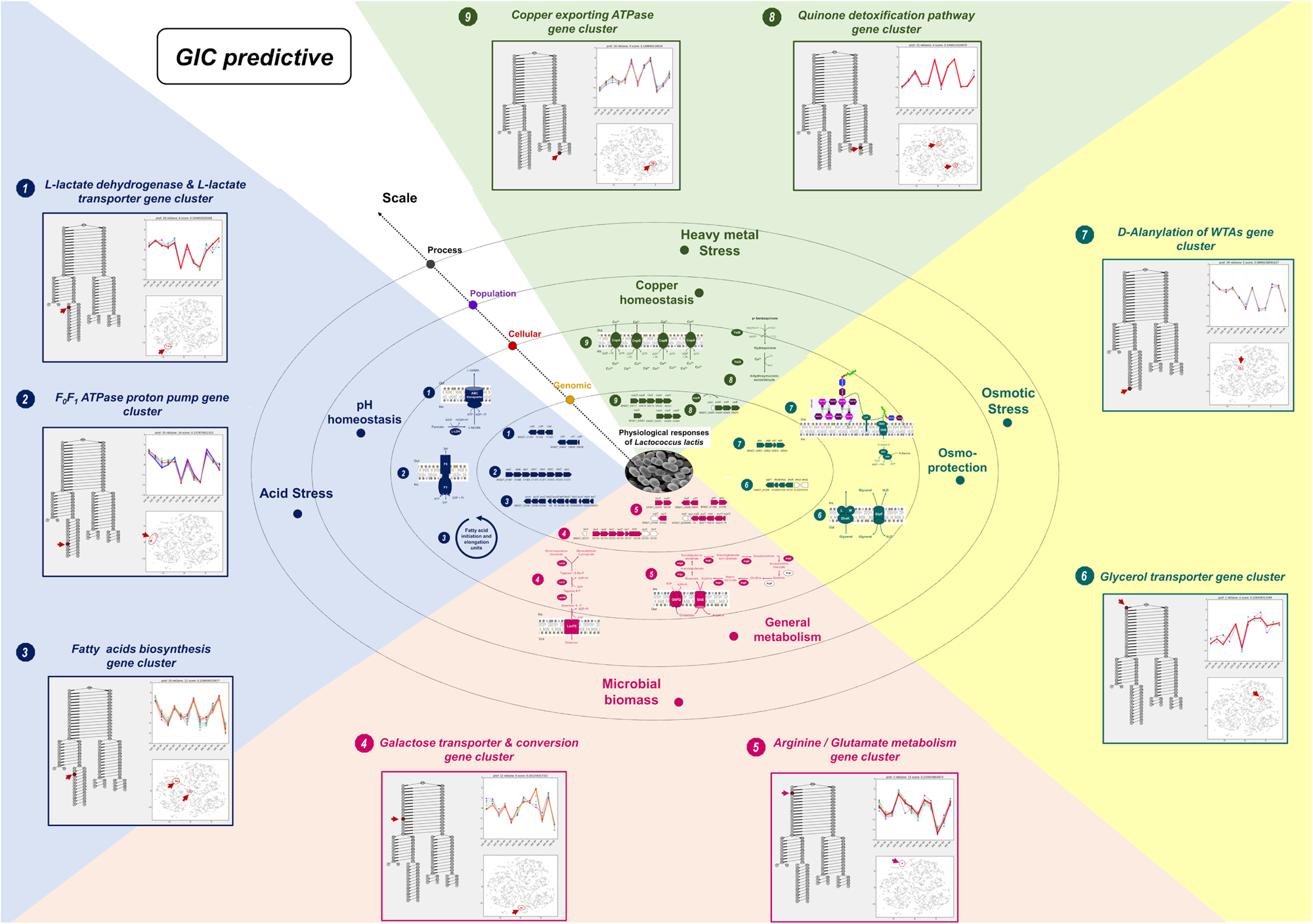
Correspondence between numerical predictions and physiological responses of the LAB to stress. The different concentric oval shapes illustrate the physiological response(s) of *Lactococcus lactis* at genomic, cellular, population and bioprocess scales, respectively. The smallest to the largest oval shape represent in order: (i) the genetic environment; (ii) the associated metabolic pathway and/or biological function; (iii) the physiological role; and (iv) metabolic responses to the bioprocess for each of the selected gene clusters highlighted in red in the dendrogram proposed by the GIC algorithm. Matches are indicated by the same color code and/or number. For the genetic environment, each gene is illustrated by a pentagon. Colors (blue, pink, cyan and green) and uncolored (white) pentagons indicate respectively the genes found and the genes missing in the cluster for the selected node. At the cellular scale, the functional protein and/or one of its subunits, which is encoded by the genes, is indicated For 1, 2, 4, 5, 6, 7 and 9 items,

In coherence with this knowledge, the equations involving the galactose genomic cluster are in almost all the predictions of the variables characterizing the cellular scale. Moreover, the genomic clusters involving *L-LDH* are proposed by this model for the prediction of 10 variables at the population scale. Under the pH-controlled *L. lactis* culture conditions, excessive L-Lactate quantities accumulation induces a strong decrease of the pH [87, 88], creating stress at the population-bioprocess level. To cope with this stress, LAB have adapted by regulating various metabolic pathways to tolerate acidifying activity and maintain physical and genetic integrity. This tolerance is ensured on the one hand, by the activity of *F0F1 ATPase proton pump* to maintain intracellular pH homeostasis. This ability to move protons out of the cell under acid conditions protects important enzymatic activities and allows long-term survival [24]. On the other hand, LAB have the ability to change the fatty acid content of their cell membrane in response to these conditions by regulating the fatty acids biosynthesis pathway [89]. This influence is seen in two variables at the cellular scale: *CFA* and *UFA* in the model. The decrease of UFAs and increase of SFAs and CFAs appears to be a bacterial strategy for maintaining cellular integrity [90]. Furthermore, this change in fatty acid composition at the end of fermentation would also improve cell survival during the freezing and freeze-drying processes [25].

In addition, lactate stress generally affects the catabolic flux of the glycolysis pathway, decreasing enzyme activity and available ATP for biomass synthesis [91], leading to unrest in cell proliferation and potential cell death. It appears in the model for *SFA* with its link to the galactose conversion and transport. Nevertheless, *Lactococci* have been able to maintain an intense level of metabolic activity by shifting the metabolism from carbohydrates to amino acids. This shift is necessary to ensure a sufficient level of ATP at the end of the stationary growth phase. Arginine, appearing in certain links, is utilized by *L. lactis* cells as a source of nitrogen which is closely linked to pyrimidine biosynthesis and also as an energy source for ATP production to meet the needs of biomass synthesis. On the other hand, it is clearly recognized that arginine also plays a role in maintaining pH homeostasis, giving tolerance to *L. lactis* cells.

Furthermore, other than its involvement in the arginine biosynthetic pathway, it is recognized that glutamate is an essential amino acid that plays a major osmoprotective role in ensuring cell viability. Other genetic elements, such as glycerol transporter and the wall teichoic acids (WTAs) D-alanylation pathway have been predicted to be equally important for maintaining cell viability. They are also present in the modeled links (Fig 8) and are essential for osmotic balance in cells. Many functional properties are related to glycerol transport (e.g., source for lipid biogenesis, facilitator of lactic acid or small solute molecules diffusion) [92], thus playing a key role in bacterial physiology. Similarly, the step of D-alanyl transfer in teichoic acid formation leading to cell wall maturation in *L. lactis* cells confers cell protection against both osmotic and acid stress [93].

Finally, during fermentation and stabilization processes (freezing and freeze-drying), LAB cells are subject to changes in the redox potential of metal ions, which are often accompanied by oxidative stress. We observe links in the model between this copper genomic cluster and all the *dtpH0.7spe* variables at the population-bioprocess scale. Among these metal ions, copper is known to be vital for the maintenance of bacterial physiology. However, excessive copper and oxidative stress can be toxic and lead to loss of viability [94]. To counteract this, LAB have developed defense mechanisms, including the use of copper transporters and detoxification systems, to maintain copper homeostasis in *L. lactis* cells (Fig 8).

Thus, based on the agreement between the physiological responses of *L. lactis* to various types of stress and the model predictions, it can be concluded that our interactive modelling tool of multi-scale biosystems is reliable, robust and useful for highlighting key physiological responses even with sparse and uncertain data. In addition, it facilitates dialogue with experts about their knowledge, even that which is not explicit.

## Discussion

The strategy used in the analysis presented here is based on a structured combination of automatic machine learning (or optimization) and user interactions steps. The idea was to support an intuitive human process for designing complex models, which consists in grouping partial knowledge relatively well mastered at some scales (local levels, in our scheme) and comparing them in order to progressively build a coherent global multi-scale interpretation. In this way, a dataset that is too small for efficient machine learning can still be exploited.

Artifical Intelligence (AI) is invoked (machine learning and optimization) each time a repetitive task is identified: segmentation steps in the GIC part, multilinear regression and global optimization in the IMSIE part. But the tuning of the respective roles of the machine and the human in such a context is a fairly delicate task. Computations and visualizations have been designed in a way that leave as much freedom of choice as possible to the expert user without drowning them in an ocean of information (user fatigue issues). We have proposed a balance based on initial bottom-up suggestions computed from the dataset (hierarchical segmentation for GIC, multi-linear regression for the local level and evolutionary optimization for the global level of IMSIE), yielding concrete hypotheses users can then revisit, constrain according to their own expertise, and recompute in order to build a multi-scale global interpretation. The tool was created to simplify the user’s task by organizing variables into clusters (GIC component) and giving the user the ability to specify constraints on groups of variables, making it convenient for a multi-scale context. We opted to present local and global models as graphical networks to better communicate key quality measurements such as precision, complexity, and other statistics. The edge colors in the graph provide an overview of the measurements and allow for quick identification of salient sub-graphs (very good or very bad).

We have seen in the use-case described above that the back and forth between local and global levels, user interactions and computations, allowed the efficient testing and questioning of various plausible hypotheses emerging from the examination of the available dataset. It has been convenient for instance to let variables of the cellular and population-bioprocess scales depend on variables at the genomic scale, and equally easy to prohibit their dependence on temperature and growth phase.

In the use-case (Production and Freeze-drying process of Lactic Acid Bacteria), the proposed interactive process has been proved useful for:

- **Retrieving known facts**, which allows the user to build trust in the system. *Lactococcus lactis* is a well-known system, so that the final graph was considered as representing a biological reality under the experimental conditions of the data. This model was also expected to help quantify some known facts on the available dataset: for the current experiment, the user estimated the shape of the equations and their parameters, which was interesting to quantify the relative importance of the variables inside an equation, but the absolute values were not considered as directly interpretable. However, developing additional visualizations to explore the behaviour of an equation with respect to the data and its uncertainties was considered interesting.
- **Integrating the different interpretations into a global model (new knowledge) and confront it with the dataset**. The graph enables correlation of different scales that provide insight into the global response of the microorganism under cell preparation conditions. It shows the effect of the process on bacterial physiology at the cellular level (membranes) and how it is regulated at the molecular level through the clusters highlighted by GIC.
- **Summarizing and simplifying the current knowledge on the process**: from an initial model with 2,741 genes, 33 variables and 165 equations to a synthetic version with 11 gene clusters, 16 variables associated to only 16 equations.

Of course, regarding the size of the dataset, the resulting model is more a combination of relevant and coherent assumptions at different scales (with respect to the dataset and the current expertise) than as a “digital twin” able to precisely mimic and predict the system of interest. While most of these hypotheses simply confirm expert insights, some of them reveal coherent hypotheses that have the potential to trigger new experimental studies.

## Conclusions

The aim of the use-case outlined in this paper was to assess an interactive modelling scheme in real-world conditions, making use of both a real-world dataset and expert input. The use-case that was considered is related to the multi-scale engineering process of freeze-drying of Lactic Acid Bacteria, which presents several challenges including: (i) the paucity and variability of the experimental dataset, (ii) the complexity and multi-scale nature of biological phenomena, and (iii) the wide knowledge about the biological mechanisms involved in this process that are not captured in the data (experts knowledge). In such a case it is obvious that classical machine learning approaches fail. We have proposed a way to exploit the power of machine learning techniques while letting domain experts conveniently control the modelling process and inject their insights using adapted visualizations and user interactions.

The experiment proved that it was possible for an expert to quickly identify a relevant group of genes, then elaborate a series of coherent local models, to finally building a global model that reasonably fits the dataset. A domain expert was also able to test the overall consistency of various hypotheses and compare them to the available data. The resulting multi-scale model features the experts’ current understanding of the process and reveals interesting hypotheses which can then be explored in future experiments. To the best of our knowledge, we produced the first multi-scale model that represents the Lactic Acid Bacteria freeze-drying process.

This interactive modelling scheme, based on the iterative use of GIC and IMSIE, can be transferred to other strains of Lactic Acid Bacteria with other experimental settings. It can also be applied to other multi-scale biological systems. This will require the integration of other types of models like Bayesian network [95], which are may be more suitable to characterize uncertain relationships between variables. Systems of differential equations may also be considered to deal with time series data. Visualization may also be improved, for instance by displaying various types of statistical information, say to identify outliers or to uncertainty sources (measurement error, model parameter sensitivity, etc). User interactions may be extended to let domain experts edit equations and models or create new variables.

The user study described in this paper demonstrates the viability of the proposed approach: the experiment was conducted using a well-known bacteria and a challenging dataset that is too sparse and ill-conditioned for classical machine learning. The experiment successfully showed that combining the experimental data with expert knowledge can help form a comprehensive representation (or a simplified model). Three novel features have been particularly valued by our domain experts: (i) the multi-scale integration as traditional tools like KEGG [96] are very efficient for representing metabolic pathways but lack the capability to integrate variables at different scales; (ii) the precision/complexity trade-off (Pareto fronts) made visible as ordered lists of models; (iii) the ability to switch between local and global modelling perspectives, as all representations can be revisited as needed.

The interactivity of the proposed approach was identified by our domain expert as a clear benefit, and the multiple scale perspectives were seen as a potential solution for reconciling contradictory experimental interpretations. Further experiments will be conducted using different biological systems and datasets to fully evaluate the capabilities of our approach.

## Acknowledgments

We aknowledge Eric Dugat-Bony (Université Paris-Saclay, AgroParisTech, INRAE, UMR SayFood) for his expertise on omics data, Thomas Chabin (Université Paris-Saclay, AgroParisTech, INRAE, UMR GMPA) and Maha Bouzaiene (ISC-PIF, CNRS) for their contributions to algorithmic developments, Jean-Daniel Fekete (INRIA, AVIZ Team) for his advises and guidance during the design process of **Biosys-LiDeOGraM** and Hélène Velly for providing the data of her PhD.

## Acronyms

AI: Artifical Intelligence.
CFA: cyclic fatty acids.
DB-SCAN: density-based spatial clustering of applications with noise.
DE: differentially expressed.
ES: early stationary growth-phase.
GIC: Genomic Interactive Clustering.
IMSIE: Interactive Multi-Scale modellIng Exploration.
LAB: Lactic Acid Bacteria.
LS: late stationary growth-phase.
SFA: saturated fatty acids.
t-SNE: t-distributed stochastic neighbor embedding.
UFA: unsaturated fatty acids.
UFC: colony-forming units.
VA: Visual Analytics.

